# A multiplex extracellular interactome screening method employing high-avidity nanoparticles

**DOI:** 10.1101/2025.07.09.663943

**Authors:** Michael A. Anaya, Maxine L. Wang, Elisa Gonzalez, Annie W. Lam, Pavee Vasnarungruengkul, Jost Vielmetter, Woj M. Wojtowicz, Kai Zinn

## Abstract

Metazoan cells signal to each other via direct contact between cell surface proteins (CSPs) and by interactions of CSP receptors with secreted ligands. CSP extracellular domain (ECD) interactions control organ development and physiology and are perturbed in disease states. However, because they cannot be accurately assessed using standard high-throughput screening techniques, they are underrepresented in protein interaction databases. Many ECD interactions are of low affinity, and their detection *in vitro* requires taking advantage of avidity effects, typically by multimerization of fusion proteins. Assays that test only one or a few interactions in each binding reaction are inadequate for global interactome screening. Here we describe a new multiplex method that uses purified dimeric ECD fusion proteins coupled to 60-mer nanoparticles as soluble prey, and the same dimers coupled to spectrally distinguishable fluorescent microspheres (beads) as bait. We add one prey to a mixture of up to 500 baits in a single well, then use a Luminex FLEXMAP 3D (FM3D) instrument to read out bait identity and prey binding. The FM3D measures the fluorescent dye ratio for each bead and simultaneously determines the amount of epitope-tagged prey bound to that bead. We use the method, denoted as the Multiplex Interactome Assay (MPIA), to analyze a proof-of-concept (PoC) set of 41 CSPs and secreted protens that is derived from larger collections examined in two interactome screens that used ELISA-based binding assays. By analyzing interactions among PoC proteins, we compared the MPIA with earlier screening methods. The MPIA has a dynamic range that is at least 30-fold greater than ELISA-based assays and appears to be more sensitive. By coupling the MPIA to an automated protein production and purification platform, we hope to be able to conduct a screen for interactions among thousands of human CSPs and secreted ligands.

## Introduction

Cell surface interactions are central to all aspects of metazoan physiology, because they define whether and how cells communicate with each other. The interactions that occur at the extracellular side of the cell’s plasma membrane, the biological interface between cells and between a cell and its environment, are critical for a myriad of processes, including organ development, maintaining homeostasis within organs, and controlling the immune system’s responses to infectious agents and tissue damage. Dysregulation of these cell surface interactions is central to disease processes.

Communication between cells can occur via cell-cell contact mediated by cell surface proteins (CSPs) or by binding of secreted ligands to CSPs. Knowledge of human CSP interaction partners is essential for assessing therapeutic potential. For example, PD-1, a negative regulator of T cell function, was discovered in 1992, but its therapeutic value only became clear when its ligand PD-L1 was identified a decade later and found to be expressed on tumor cells. Anti-PD-1 and anti-PD-L1 antibody drugs are now widely used for immunotherapy and can induce long-term remission for some lethal metastatic cancers (reviewed by (Ishida 2020; Parvez et al. 2023)).

Multiple high-throughput methods have been developed that allow large-scale screening of protein-protein interactions (PPIs). Large-scale screens have been performed for several metazoan species and thousands of PPIs have been identified. Despite this progress, the interaction networks of CSP extracellular domains (ECDs) are still poorly defined, because these PPIs cannot be accurately assessed using the yeast two-hybrid screening assay (Y2H) or affinity purification/mass spectrometry (AP-MS), which are the most common methods for interactome screening. ECDs and secreted proteins are usually glycosylated and have disulfide bonds, and their expression requires trafficking through the endoplasmic reticulum and Golgi, making them incompatible with standard Y2H methods. Interactions among CSP ECDs are often weak, with K_D_s in the µM range, making them difficult to capture using AP-MS due to their short lifetimes (reviewed by (Shilts and Wright 2024)). Many ECD interactions detected by these high-throughput assays are listed in databases such as BioGrid, BioPlex, STRING, and IntAct, but most of these are indirect interactions or false positives.

CSP-CSP interactions, even those with low affinities, can create stable adhesive interactions between cells through avidity effects. There are often many copies of each protein packed densely at cell interfaces, so the local concentration is high. Researchers have taken advantage of avidity through ECD clustering to detect CSP interactions *in vitro*. Avidity enhancement can be achieved by incorporating multimerization domains (Capon et al. 1989; Tomschy et al. 1996) into soluble ECD fusion proteins and creating dense arrays of ECDs on surfaces. Experimental platforms that utilize clustering or multimerization include ELISA-based binding assays (Bushell et al. 2008; Wojtowicz et al. 2007; Özkan et al. 2013), protein microarrays (Sun et al. 2012), cell-signaling assays (Barrow et al. 2018), cell-surface staining of reverse-transfected cell microarrays (Turner et al. 2013), and bead-based assays (Husain et al. 2019; Li et al. 2017). ELISA-based assays, which are easy to perform and have been reported to have a low false positive rate, are the most commonly used methods for high-throughput cell-surface interactome screening (Bushell et al. 2008; Wojtowicz et al. 2007; Özkan et al. 2013; Visser et al. 2015; Ranaivoson et al. 2019; Verschueren et al. 2020; Wojtowicz et al. 2020; Shilts et al. 2022). The foundation for the work in this paper was provided by a semi-automated ELISA-based interactome (ECIA) screen of 564 human ECDs conducted by groups at Stanford and Caltech. These included all non-antibody, non T-cell receptor immunoglobulin superfamily (IgSF) proteins, as well as some proteins with other domains. IgSF proteins are divided into many subfamilies, and IgSF is the most highly represented protein domain in CSP ECDs in all metazoans. This screen, the Human Interactome Project 1 (HIP1), involved assessing 318,096 pairwise interactions (Wojtowicz et al. 2020). We are now conducting a screen of >2200 human CSPs and secreted ligands (∼5 million pairwise interactions), which is beyond the capacity of ELISA-based screening methods. To allow us to conduct such a screen, we need a method in which many interactions can be assessed in every well. In 2017, we developed a multiplexed method, the Bio-Plex Interactome Assay (BPIA), and conducted a pilot screen of a small Drosophila protein interaction network (Li et al. 2017). The method described here evolved from that work.

The Luminex (Diasorin) system, which was licensed to Bio-Rad at the time of the 2017 screen (hence the name Bio-Plex) utilizes the principles of flow cytometry and is most commonly employed for multiplexed screening of serum samples for cytokines and autoreactive antibodies. It has also been used for multiplexed screening of protein-protein interactions (Blazer et al. 2010; Blazer et al. 2011; Rimmele et al. 2010; Li et al. 2017). The Luminex system uses magnetic polystyrene beads, called MagPlex^TM^ beads, that are embedded with different ratios of fluorescent dyes, rendering them spectrally distinct when excited by a laser. Each bead type is called a bead region. The FLEXMAP 3D (FM3D) instrument used in our experiments can distinguish up to 500 bead regions. The beads are covalently coupled to lysine groups on proteins using N-hydroxysuccinimide (NHS) chemistry. Protein-conjugated bead regions (baits) are pooled and incubated with a soluble protein (prey) to simultaneously test binding of the prey to up to 500 different bait proteins in a single well of a multi-well plate. Binding of prey to bait proteins is detected using a primary antibody against an epitope on the prey (mouse anti-V5 for our preys), followed by phycoerythrin (PE)-coupled rabbit anti-mouse secondary antibody. Streptavidin-PE can also be used if a biotinylated primary antibody is being detected. Beads flow through the FM3D in single file and are interrogated by two lasers: one to discern the identity of the bead region (which represents the identity of the bait protein), and the other to detect the PE prey signal for each bead. The machine is programmed to read 50 beads of each region before moving to the next well. The readout of the median PE signal for these beads (median fluorescence intensity, MFI) represents the amount of prey binding to bait. The assay has a high signal-to-background ratio, because strong binding of a prey to a bait bead can generate MFI readings of >30,000, vs. 30-100 for binding to control or unconjugated beads.

In standard ELISA-based screening methods, the bait and prey need to be tagged differently, so that prey can be distinguished from bait (Wojtowicz et al. 2007; Bushell et al. 2008; Özkan et al. 2013; Verschueren et al. 2020). This means that every protein needs to be expressed in two forms. In HIP1 and the BPIA pilot screen, these were fusions of ECDs to dimeric human IgG1 Fc fragments (hFc) and to pentamerized alkaline phosphatase (AP5) (Li et al. 2017; Özkan et al. 2013; Wojtowicz et al. 2020; Bushell et al. 2008). One issue with using different formats for prey and bait proteins is that they may differ in aspects like post-translational modification and aggregation, which are critical for the correct function and interaction of CSPs. Additionally, pentamerized AP5 proteins are usually expressed at lower levels than dimeric hFc proteins.

Recently, Wright and colleagues developed an improved version of their AVEXIS ELISA-based binding method, called SAVEXIS, wherein a single protein is used as both prey and bait (Shilts et al. 2022). This method employs the high-affinity biotin::streptavidin interaction to generate ECD tetramers and utilizes streptavidin to both immobilize bait protein on the plate surface and detect prey binding.

We have employed a different strategy to avoid having to make every protein in two different forms. Our method uses an adapter protein to introduce a unique epitope onto the prey. We make ECD proteins as fusions with hFc, purify them on a liquid-handling robot using an 8xHis tag, and measure their concentrations. The standard ELISA-based screening methods used ECD proteins in unpurified cell supernatants, while SAVEXIS used purified ECD proteins. Purification yields proteins with higher concentrations than in supernatants.

In two previous ELISA-based screens using unpurified supernatants, HIP1 and a screen conducted by Genentech (GNE) that tested many of the same proteins, the overlap of observed interactions between proteins that were tested in both screens was only ∼30% (Verschueren et al. 2020; Wojtowicz et al. 2020). After the start of our project, a third interactome screen containing many overlapping IgSF proteins (SAVEXIS) was published (Shilts et al. 2022), and this had a similar overlap with HIP1 and GNE. We thought that the low concordance among screens might be due to differing levels of protein expression attained by the various groups, and we hoped to address this issue by using purified proteins at known concentrations. We also wanted to make higher-avidity preys, in the hope of finding weaker interactions that might have been missed entirely in screens that test binding of pentameric prey.

Here we describe a new concept that accomplishes these goals, in which SpyTag-SpyCatcher technology (Zakeri et al. 2012) is used to covalently couple fusion proteins to beads and nanoparticles. mi3 is a virus-like nanoparticle with 60 identical subunits that can be expressed at high levels in E. coli, and SC3-mi3 has the SpyCatcher003 domain (SC3) fused to the mi3 monomer (Bruun et al. 2018; Tan et al. 2021). SpyCatcher spontaneously forms a covalent isopeptide bond with a 14 amino acid peptide called SpyTag. With the latest versions of these proteins, SpyTag003 (ST3) and SC3, this reaction occurs within 15 minutes with an efficiency of near 100%, even at concentrations as low as 10 nM (Keeble et al. 2019). This allows investigators to load SC3-mi3 particles with precisely defined ratios of two or more proteins (Cohen et al. 2021; Tan et al. 2021). In our experiments, we label a subset (∼20%) of the 60 SC3 sites on SC3-mi3 nanoparticles with a V5 epitope-tagged adapter protein, then fill the rest of the sites with an ECD-hFc-ST3 fusion protein. To generate bait, the same ECD-hFc-ST3 fusion proteins are coupled to SC3-conjugated MagPlex^TM^ beads.

In this paper, we describe the development and optimization of a screening methodology using these preys and baits, which we call the MPIA (Multiplex Interactome Assay). The screen workflow is shown in Fig. 1. To develop the assay, we screened a small “proof of concept” (PoC) set of 41 proteins, which contained 39 binding pairs found in either the HIP1 or GNE screens (Wojtowicz et al. 2020; Verschueren et al. 2020)(Fig. 2). We define the set of 39 binding pairs as the PoC network, and benchmarked the MPIA screen against HIP1 and GNE by examining its recovery of these binding pairs and its ability to find other binding pairs within the PoC set that had not been detected in earlier screens.

**Fig. 1.**
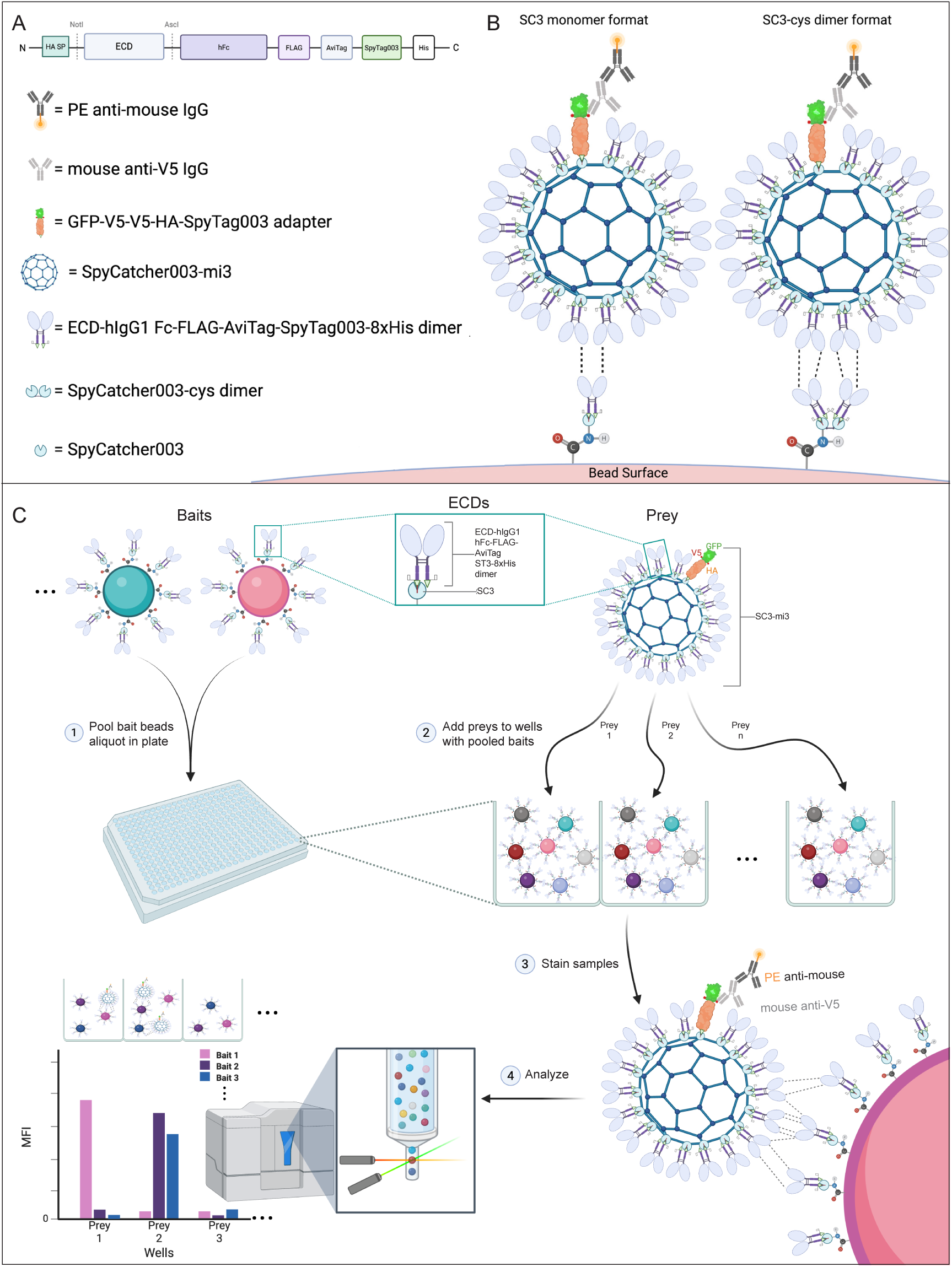
The MPIA method. A. Map of ECD constructs. ECD sequences are inserted between the Not I and Asc I sites. B. Cartoon of prey nanoparticle binding to monomeric SC3 and dimeric SC3 bait. Prey binding is detected using mouse anti-V5 and PE anti-mouse. V5 epitope on adapter indicated by red dots. C. Flowchart of the screening process. Bait beads, with each ECD coupled to a different region, are pooled and placed in wells of a plate (1). One prey is added to each well (2), and after incubation primary (anti-V5) and secondary antibody (PE) are added (3). In the FM3D, beads flow in single file past two lasers, which interrogate the bead region and read the PE signal (4). The median bead fluorescence signal (MFI) is displayed on the bar graph for three preys binding to three baits, with cartoons of the binding events above the graph. In this example, prey 1 binds to bait 1, prey 2 binds to baits 2 and 3, and prey 3 does not bind to any bait.

**Fig. 2.**
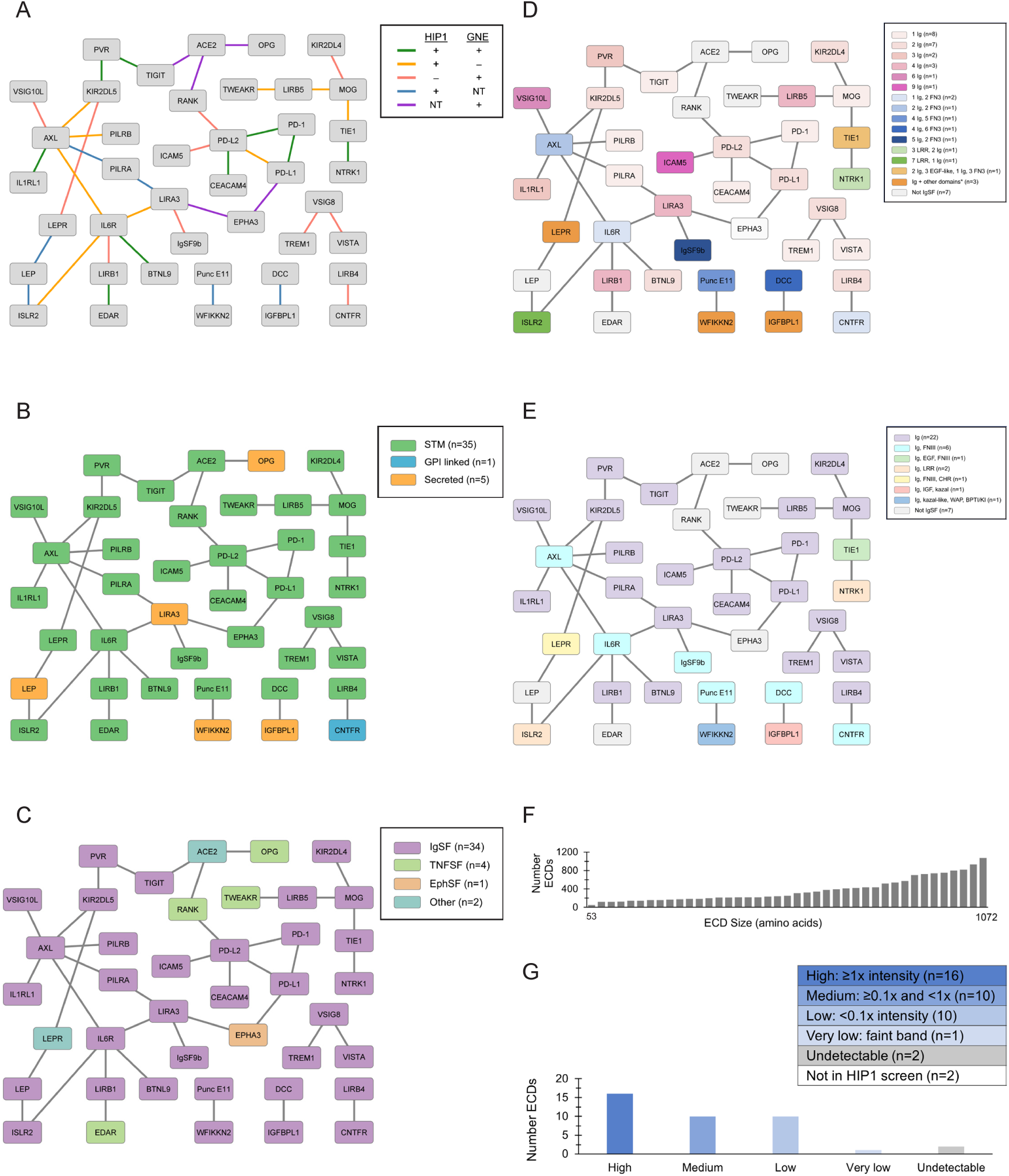
The proof of concept protein set. A. The PoC network (39 binding pairs within the PoC set). Colored lines indicate screen(s) in which binding pair was found. NT, not tested. B. Protein type. C. Protein family. D. Domain architecture in IgSF ECDs. E. Domain types in IgSF proteins. F. ECD amino acid size distribution. G. Qualitative protein expression of HIP1 ECD-Fc based on anti-His Western relative to 10 ng purified L12-Fc. Abbreviations: immunoglobulin superfamily (IgSF), tumor necrosis factor superfamily (TNFSF), ephrin superfamily (EphSF), single-pass type I transmembrane receptor (STM), glycosyl- phosphatidylinositol (GPI), Immunoglobulin (Ig), fibronectin type III (FN3), leucine-rich repeat (LRR), tumor necrosis factor receptor cysteine-rich domain (TNFR-cys), cytokine receptor homology (CHR), epidermal growth factor-like (EGF-like), whey acidic protein (WAP), bovine pancreatic trypsin inhibitor/Kunitz inhibitor (BPTI/KI). **Figure 2, data supplement (a separate file)**

We show that the MPIA method has a remarkable signal-to-background ratio (up to 1000- fold). We identified many PPIs seen in the earlier screens and also uncovered new candidate PPIs that had not been previously detected. In other work, not described here, we have also shown that each of the steps in the method can be fully automated, including production of plasmid DNA, cell transfection, protein purification, and the MPIA assay itself. We have thus developed a methodology that can allow us to execute a screen of >2200 human CSP ECDs and secreted proteins. This represents most of the genes encoding CSPs whose ECDs can be expressed in a soluble form, separate from the plasma membrane, as well as genes encoding orphan secreted proteins and secreted proteins that we anticipate may have additional unknown receptors. We estimate that, when the proteins have been expressed and purified, 2200 preys could be constructed and screened against 2200 baits within a few months using this technology.

## Results

### Selection of the PoC protein set

We used multiple criteria to select a PoC PPI network from the combined HIP1 and GNE screens (Wojtowicz et al. 2020; Verschueren et al. 2020) for validating and benchmarking the MPIA as a higher-throughput alternative to the standard ELISA-based screening methods (Fig. 2A and Fig. 2, data supplement). The PoC protein set has 41 proteins, and the PoC network defined by HIP and GNE comprises 39 binding pairs within this set. Of these 39, some were tested in both HIP1 and GNE while others were tested in only one (see below). Of course, there are many other possible binding pairs within the PoC protein set that were not observed by either HIP or GNE. The PoC network contains only high-confidence PPIs from the HIP1 and GNE screens, defined as those with fold-over-background (FOB) values of ≥5 in the HIP1 screen and as POS-HICONF in the GNE screen. We tested each binding pair in both orientations (prey 1::bait 2 and prey 2::bait 1). Since each binding pair corresponds to two possible PPIs, the PoC network of 39 binding pairs examined in our screen comprises 78 unique PPIs. However, HIP1 only tested 68 of these, and GNE tested 61. The pattern of binding pairs (i.e., lines connecting proteins in the network) defines the interactome map, while detection of PPIs in both orientations is important for evaluating the robustness of the method. Bidirectional detection of PPIs increases confidence in the identification of a binding pair, but ELISA-based screening methods often observe binding in only one orientation (unidirectional). Binding pairs and PPIs are both important metrics for validating and benchmarking the MPIA.

The PoC proteins were selected to maximize our ability to assess both metrics, while simultaneously ensuring that a diverse set of proteins was included. We included three major types of proteins: single transmembrane (STM)(n=35), secreted (n=5), and GPI-linked (n=1)(Fig. 2B). We selected proteins covering a broad ECD size range (53-1072 amino acids) (Fig. 2F) and belonging to different families, including IgSF (n=34) and tumor necrosis factor (TNFSF, n=4), as well as one ephrin receptor and two proteins that contain other miscellaneous domains (peptidase M2, collectrin-like, 4-helix bundle) (Fig. 2C). Furthermore, we selected several IgSF proteins that contain additional domain types (e.g., EGF-like, Kazal-like, leucine-rich repeats (LRR), fibronectin type III (FN3)) (Fig. 2D) and varying domain architectures (e.g., different numbers of Ig domains, different numbers and combinations of Ig and other domains) (Fig. 2E). Lastly, the ECD-hFc-ST3 proteins used in the MPIA contain different C-terminal tags from the ECD-Fc proteins used in our HIP1 screen. To examine whether the altered C-terminal region in ECD-hFc-ST3 proteins has an effect on protein expression levels, we included ECDs that exhibited a broad distribution of expression levels in the HIP1 screen (Fig. 2G). Of the 41 proteins selected for the PoC set, 39 were tested in HIP1 (i.e., two proteins were screened only by GNE).

To benchmark the MPIA against the standard ELISA-based methods, we considered both the identity of the binding pairs (connecting lines in the network) and the PPIs (directionality) observed in the HIP1 and GNE screens. We included binding pairs that were observed in both the HIP1 and GNE screens (n=9), pairs tested in both the HIP1 and GNE screens but only observed in one screen (n=19), and pairs tested in only one of the two screens (n=11) (Fig. 2A). Five of the GNE binding pairs were only tested in one prey::bait orientation. We additionally included binding pairs with PPIs observed bidirectionally and binding pairs with PPIs observed only unidirectionally in one or both screens. Our rationale for including binding pairs tested in both screens but only seen in one screen and binding pairs observed only unidirectionally was to assess whether the MPIA provides higher sensitivity, resulting in detection of PPIs that were false negatives in one of the two ELISA-based screens. Higher sensitivity in the MPIA might arise from an increase in avidity of the prey SC3-mi3 nanoparticles and/or from higher levels of ECD on the bait beads and prey nanoparticles as compared with the HIP1 and GNE screens, which used proteins in unpurified supernatants.

The overlap of binding partners observed in three large interactome screens of cell-surface and secreted proteins using ELISA-based binding methods (HIP1, GNE, and SAVEXIS) was about 30%. Although this number seems low, in their recent review, Wright and colleagues emphasized that the ∼30% overlap in these three systematic empirical datasets is much higher than the overlap of ECD binding partners reported across a majority of six widely used databases of protein-protein interactions (3%) (Shilts and Wright 2024). The low overlap among these databases likely arises due to the prevalence of false positive PPIs for ECDs in large Y2H and AP-MS datasets, as discussed above. By contrast, the false positive rate of PPIs observed in the three ELISA-based screens is argued to be lower because most randomly-selected PPIs exhibited binding in orthogonal assays such as surface plasmon resonance (SPR) and cell binding (Wojtowicz et al. 2020; Verschueren et al. 2020; Shilts et al. 2022). As such, the ∼30% overlap between screens is likely due primarily to false negatives (i.e., a PPI observed in one screen is not observed in other screens). It is still unclear why the false negative rate is so high. In developing our method, we imposed the metric that detection of >30% of the PPIs in the PoC network was necessary to validate the MPIA as a higher-throughput alternative to ELISA-based screening.

### Generation of ECD fusion proteins

To express the PoC proteins in an appropriate form for the MPIA, we first subcloned DNA encoding the ECDs from plasmids used for protein expression in HIP1 into a new vector, which inserted the ECD into a framework where proteins are fused on the N-terminal end to the influenza hemagglutinin (HA) signal peptide (SP) sequence, and on the C-terminal end to hFc-FLAG-AviTag-ST3-8xHis (fusion proteins are abbreviated ECD-hFc-ST3). We tested a representative set of proteins with other SPs but found that none consistently yielded higher levels of protein than the HA SP (data not shown), which is the same SP used in HIP1. Each protein was expressed in a 30 ml Expi293 suspension culture and purified from cell supernatants using Ni-NTA resin to generate a final preparation of 5 ml, which represents a 6-fold increase in concentration relative to the supernatant. At least three preparations were made for each protein and protein concentrations were determined using densitometry of bands on Coomassie Blue-stained gels. The PoC proteins were present in the purified preparations at widely varying levels, ranging from 100 nM (the lowest concentration we could quantitate via densitometry) to 60 µM (Fig. 3, data supplement). Two to four proteins were present at <100 nM in our preparations. These proteins were still detectable by Western blotting.

**Fig. 3.**
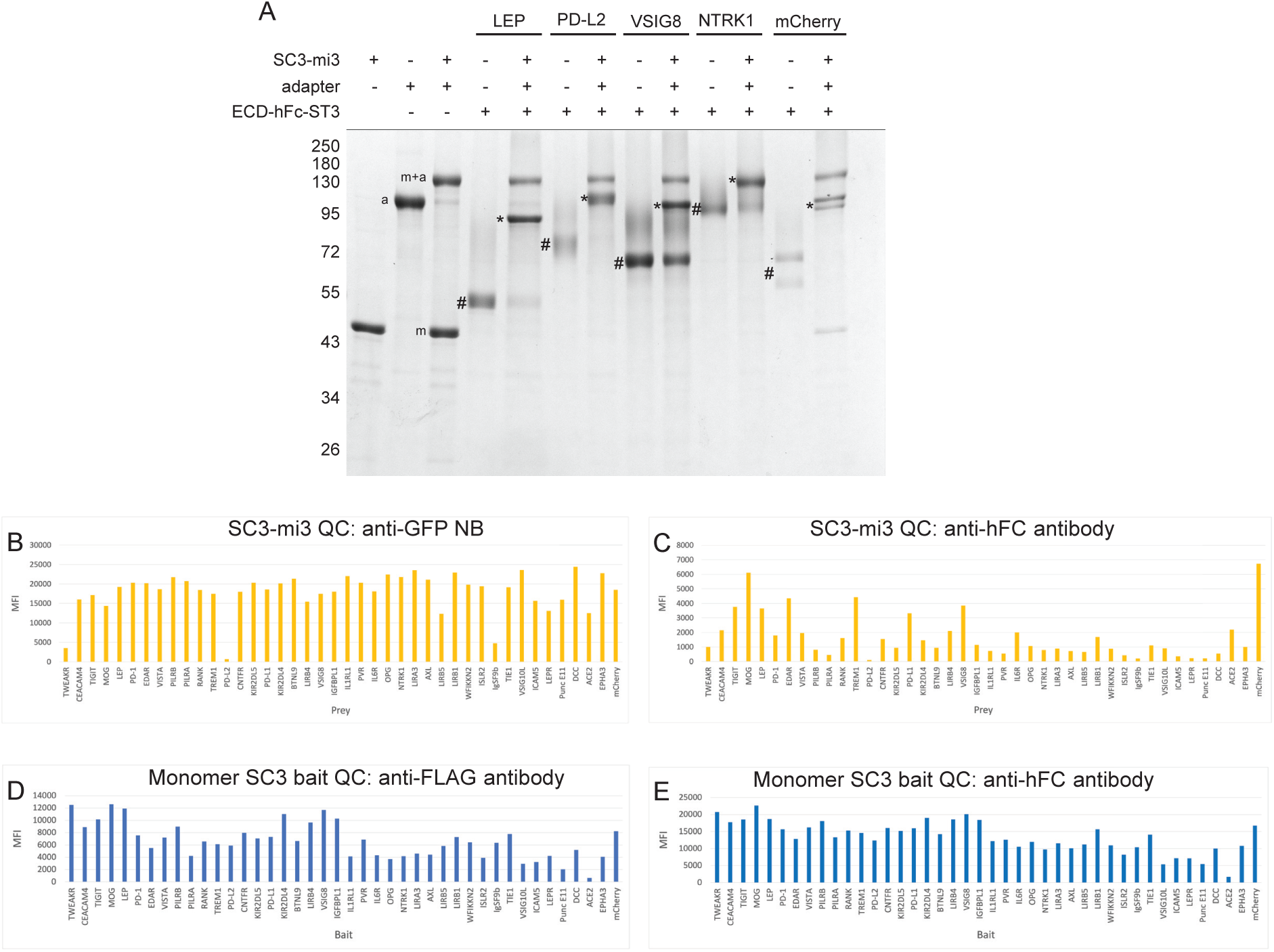
Generation and quality control of prey nanoparticles and bait beads. A. SDS-PAGE analysis of adapter coupling to SC3-mi3, and coupling of four different ECD-hFc-ST3 proteins to SC3-mi3-adapter complexes. m, uncoupled SC3-mi3 monomer band (this band is depleted when all SC3 sites are coupled to excess ECD, so it is absent except in the mCherry lane, where less ECD was added); a, GFP-V5-HA-ST3 adapter; m+a, adapter covalently linked to SC3-mi3 (this band is present in all lanes in which ECD is coupled to the SC3-mi3-adapter complex); #, free ECD-hFc-ST3 protein (since ECD was in excess, these bands are also visible in lanes containing SC3-mi3-adapter-ECD complexes); *ECD-hFc-ST3 covalently linked to SC3-mi3 (this band runs on top of the SC3-mi3-adapter band for NTRK1). Uncoupled and coupled mCherry-hFc-ST3 are doublet bands. Marker protein band positions indicated, in kD. B. Bar graph of MFI values for PoC prey nanoparticle binding to beads coupled to anti-GFP NB. In B-E, proteins are ordered from smallest to largest ECD. PD-L2 gave a low signal in this experiment, for unknown reasons. However, this PD-L2 prey still bound to PD-1 and CEACAM4. C. Bar graph of MFI values for PoC prey nanoparticle binding to beads coupled to anti-hFc antibodies. Note that these are highly variable, and that some preys with high anti-GFP signals (e.g., PVR) have very low anti-Fc signals, suggesting steric hindrance in access to the Fc portion of the fusion protein when it is coupled to the nanoparticle. D. Bar graph of MFI values for anti-FLAG prey binding to PoC bait beads. E. Bar graph of MFI values for anti-hFc prey binding to PoC bait beads. Note that the anti-FLAG and anti-Fc graphs have similar profiles. PVR bait anti-Fc signals are much higher than for prey, suggesting that there is less steric hindrance for protein on beads. Protein on beads is probably present at a much lower density than on the nanoparticle. **Figure 3, data supplement (a separate file)**

### Development of the SC3-mi3 nanoparticle prey system

In the MPIA, the same ECD-hFc-ST3 protein is coupled to prey SC3-mi3 nanoparticles (Tan et al. 2021) and to bait beads. To detect binding of prey to bait, we couple an adapter protein with a unique epitope (a V5 tag) to a fraction of the SC3 sites on SC3-mi3 nanoparticles. The V5 epitope on the adapter allows detection of prey binding to bait using a mouse anti-V5 antibody, followed by PE-conjugated goat anti-mouse antibody. When a fraction of the SC3 sites were occupied by the adapter and the rest were filled with ECD-hFc-ST3 protein containing a large ECD, we observed that N-terminal V5 tags on short adapters (e.g., GFP fusion proteins) were not detected well by anti-V5 antibodies (data not shown). Accordingly, we made an adapter using the influenza HA protein, which is a long, rigid trimeric rod that is 13.5 nm in length. After filling all SC3 sites with ECD-hFc-ST3s, this adapter was accessible to anti-V5 antibodies even when large ECDs were used. The HA adapter has the structure GFP-2xV5-HA-ST3-8xHis (abbreviated GFP-V5- HA-ST3).

To generate prey particles, we first coupled GFP-V5-HA-ST3 to a subset of the 60 SC3 sites on SC3-mi3. We found that loading of ∼20% of the SC3 sites with adapter was optimal to obtain maximal V5 signals for prey::bait binding (data not shown). On an SDS-PAGE gel, the 20% filled SC3-mi3-adapter complex runs as two major bands: SC3-mi3-adapter and SC3-mi3 monomer (Fig. 3A). Following adapter coupling, we incubated SC3-mi3-adapter nanoparticles with ECD-hFc-ST3s using a ratio of ∼1.5-fold over the remaining SC3-mi3 sites to ensure that all SC3 sites were filled. Covalent coupling of ECD-hFc-ST3 to SC3-mi3 increases the molecular weight of the SC3-mi3 subunit, which can be visualized by SDS-PAGE. After filling the remaining SC3 sites on the SC3-mi3-adapter with an ECD, two additional bands are observed: 1) SC3-mi3- adapter-ECD and 2) uncoupled excess ECD-hFc-ST3. The ECD-hFc-ST3 band is reduced in intensity relative to the amount of input protein. If the SC3 sites on SC3-mi3 are completely occupied by ECD-hFc-ST3, the SC3-mi3 monomer band disappears (Fig. 3A).

When increasing concentrations of SC3-mi3-adapter-ECD were incubated with bait beads, we observed that background binding to bait beads was not affected by prey concentration when optimal blocking buffers were used (see below), so we were able to use preys at high concentrations. For proteins expressed at sufficiently high levels, we used preys at a concentration of 300 nM ECD. Since SC3-mi3-adapter-ECD prey particles had to be diluted 3- fold into blocking buffer for the binding reaction, we made preys using 1.5 µM ECD (0.75 µM of ECD-hFc-ST3 dimer) in the coupling reaction. For purified proteins with concentrations lower than 1.5 µM (between nine and 14 proteins depending upon the purified preparation; Fig. 3, data supplement), we mixed undiluted ECD-hFc-ST3 protein with a reduced amount of SC3-mi3-adapter that would allow for filling of all SC3 sites. In the binding reaction with the beads, these lower concentration preys were thus present at ECD concentrations well below 300 nM.

For every preparation of prey particles, we employed the FM3D to measure SC3-mi3 nanoparticle concentration and confirm loading of ECD onto the nanoparticle using two quality control (QC) beads. These beads were coupled to anti-GFP nanobody (NB) or goat anti-hFc antibody using NHS chemistry. The anti-GFP NB and anti-hFc antibody bind the GFP-V5-HA-ST3 adapter and ECD-hFc-ST3 on the prey, respectively. Binding of prey to the beads via the NB or antibody is detected using mouse anti-V5 antibody, followed by PE-conjugated goat anti-mouse antibody. Figs. 3B-C show bar graphs of the MFI signals for the preys with QC beads. We observed consistent MFI signals for anti-GFP NB beads, whereas anti-hFc beads generated highly variable signals (Figs. 3B-C). This is likely because the N-terminus of the rigid GFP-V5- HA-ST3 adapter protrudes beyond the “envelope” formed by ECDs. In contrast, the anti-hFc antibody contacts epitopes that are C-terminal to the ECD and may therefore vary in accessibility from one ECD to another due to steric hindrance, affecting the ability of the prey particles to bind to anti-hFc beads. Thus, the purpose of the anti-hFc QC is to establish that SC3-mi3 particles contain ECDs, not to measure the amount of ECD on each particle. To evaluate the amount of ECD-hFc-ST3 coupled to SC3-mi3-adapter, we analyzed prey using SDS-PAGE, as in Fig. 3A. However, since it would be impractical to run gels for every prey nanoparticle in a large screen comprising hundreds of preys, QC on the FM3D provides a high-throughput way to confirm that coupling of ECD-hFc-ST3 to SC3-mi3 occurred. The ST3-SC3 reaction is robust and highly reproducible, so we expect that the reaction will go to completion and that the nanoparticles will be completely filled with ECD if excess ECD is used.

### Generation of bait beads

To make bait beads, we employed a strategy in which SC3 is directly coupled to beads using NHS chemistry, followed by immobilization of ECD-hFc-ST3 bait proteins onto the SC3-coupled beads. Such a strategy confers multiple advantages. First, it allowed us to directly couple only a single “capture” protein, namely SC3, to all of the bead regions. Second, by immobilizing ECD-hFc-ST3 onto SC3-coupled beads rather than directly coupling each ECD-hFc-ST3 to beads using NHS chemistry, we ensure that the ECDs are displayed on the beads without chemical modification of lysines. Third, because ST3 is at the C-terminus of ECD-hFc-ST3, proteins are oriented on the bead with the ECD facing outward. Lastly, the concentration of ECD-hFc-ST3 needed to saturate SC3-coupled beads is lower than what would be needed for direct coupling, enabling us to saturate beads even with poorly expressed proteins.

Following SC3 coupling to beads, beads were loaded with ECD-hFc-ST3. We found that between 100 nM and 500 nM was required to achieve maximum bead loading (data not shown), so we used 500 nM ECD-hFc-ST3 (250 µM ECD-hFc-ST3 dimer) for proteins expressed at sufficiently high levels. For ECD-hFc-ST3 proteins with concentrations less than 500 nM (seven and 10 proteins in the first and second purifications, respectively), we added undiluted protein to the beads. As described below, we later modified the SC3 beads so that we were able to attain high levels of ECD-hFc-ST3 on the beads even for proteins at concentrations as low as 10 nM.

To confirm that ECD-hFc-ST3 was loaded onto the bait beads, we stained the beads with mouse anti-hFc and mouse anti-FLAG antibodies (ECD-hFc-ST3 contains a FLAG epitope), followed by PE-conjugated goat anti-mouse secondary antibody, and analyzed the beads on the FM3D. We observed variability among signals for bait beads loaded with the same ECD concentrations (Figs. 3D-E). Presumably this is because both the hFc and FLAG epitopes are C-terminal to the ECD and thus closer to the bead, so steric hindrance affects how much primary antibody can bind to the ECD-hFc-ST3. Because of this, as with the prey QC (Fig. 3B-C), the purpose of the bait bead QC is to confirm that ECD-hFc-ST3 loading onto the bead was successful, not to provide quantitative information.

After each ECD-hFc-ST3 bait protein was coupled to an assigned bead region, the beads were pooled to create the multiplexed bait bead set. At this point, any remaining uncoupled SC3 sites on the beads were blocked by incubation with synthetic ST3 peptide, followed by additional washes and final storage in blocking buffer.

### Screening the PoC proteins

To run a screen, we added 500 beads for each of the baits to wells of a 96-well plate. We also included two bead regions as controls for nonspecific binding of prey to bait beads. One contained no ECD-hFc-ST3 (SC3-only, blocked with ST3 peptide) and the other was loaded with mCherry- hFc-ST3. SC3-mi3-adapter-ECD preys (at 300 nM ECD where possible, or less for poorly expressed proteins) were added to the multiplexed bait bead set. We also included the control prey SC3-mi3-adapter-mCherry. Following incubation of prey with bait beads for 2 hr at room temperature, beads were sequentially incubated with mouse anti-V5 and PE goat anti-mouse antibodies to detect the prey bound to the beads. In separate control wells that did not contain prey, we added mouse anti-hFc and anti-FLAG, which detect ECD-hFc-ST3 on the multiplexed bait bead set. Following antibody incubation and washing, beads were analyzed on the FM3D, which records MFIs for each bead region.

We did an extensive series of tests of blocking buffers to determine conditions that minimize nonspecific binding of prey to control beads. Remarkably, we observed that when 1% casein was used as a blocking agent, nonspecific binding signals were >10-fold lower than with any other buffer (data not shown). Using these conditions, we conducted screens with the PoC set proteins that test pairwise binding of every combination of the 41 proteins (1,681 prey-bait pairs). The screen results from two runs (M1 and M2) with different purified protein preparations are shown in the graphs of Figs. 4A-B, which are histograms of binned MFI values (raw MFI data in Fig. 6, data supplement). The vast majority of data points have very low MFI signals (<100), while a small number have MFI values ranging up to >20,000. We also show Empirical Cumulative Distribution (ECDF) plots in which MFI values from lowest to highest are on the x-axis, and their percentiles are on the y-axis (Figs. 4C-D). The ECDF curves are non-normally distributed, with >95% of the data points on the steep ascending portion with low MFI values, then a point of maximum curvature (knee) where the slope transitions from a steep to a gentle slope, and a small number of points on the right side of the knee, representing the minority population. The ECDFs indicate that most prey-bait combinations (non-hits) do not result in any specific binding events and thus generate low MFI signals to the left of the knee, comparable to mCherry preys with any bait, or any prey with mCherry or ST3-only beads. In two separate experiments run with different ECD-hFc-ST3 protein preparations, the knees, which may represent the border between hits and non-hits for a particular run, are at MFIs of 930 and 830. The locations of the knees differ more for repeat experiments conducted on different days than for duplicate samples run on the same day (see below), because the readout is the signal following primary and secondary antibody staining.

**Fig. 4.**
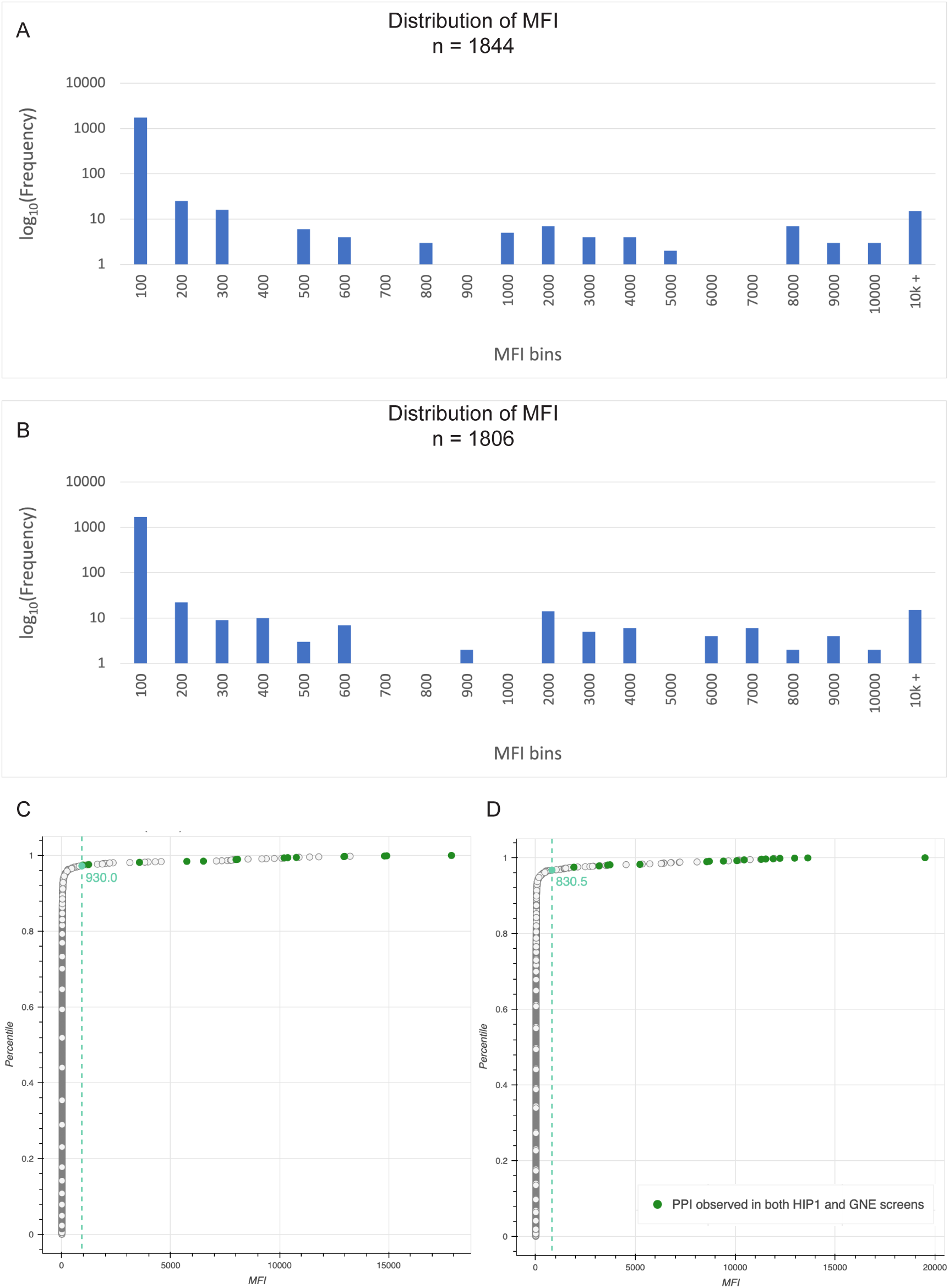
Screening the PoC set proteins using the MPIA. A. Histogram of binned MFI values, for run M1. The y-axis is on a log scale, and the vast majority of data points have MFIs of <100. Note that there is a gap in the distribution around 700-900 for both runs. This correlates well with the ECDF knee position. B. Histogram of binned MFI values for run M2. C. ECDF plot of run M1. The x-axis is MFI and the y-axis is the cumulative probability. For a given value on the x-axis, the y-axis value represents the fraction of data points that have a value less than or equal to the x-axis value. For example, the data point (550, 0.965) means that 96.5% of measured MFIs have a value less than or equal to 550. The 17 green dots are the points representing hits that were also found in both the HIP1 and GNE screens. Note that all of these are to the right of the point of maximum curvature (knee), indicated by a teal dot and dotted line, where the curve changes from the steep region where most values occur to the gentle region that encompasses the rare hits. The knee is at MFI 930. D. ECDF plot of run M2, with common (MPIA+ HIP+ GNE+) hits and knee point indicated as in C. The knee is at MFI 831.

We first evaluated the results of these screens relative to the binding pairs seen in *both* the HIP1 and GNE screens. We found that we were able to detect 100% of these binding pairs (9/9). Each pair has two PPIs, and we found 94% of the PPIs that comprise them (17/18). HIP1 and GNE each found 14 of these 18 PPIs (see Fig. 4, data supplement). On the ECDF plots, these 17 PPIs corresponding to the HIP1+GNE binding pairs are all to the right of the knee (green dots in Figs. 4C-D). However, the MFIs for PPIs defining binding pairs that were seen in only one of the earlier screens were often to the left of the knee. Also, PPIs involving proteins expressed at low levels tended to have low MFIs. This suggested that it would be valuable to try to increase the sensitivity of the MPIA. We did this by increasing the density of ECD on the beads, as described below.

### Dimeric SC3 beads generate stronger signals than monomeric beads

Because some of our proteins were expressed at low levels, we were concerned that we would miss some interactions that might have been detected if there were more ECD on the bait bead. To address this issue, we sought to increase ECD loading, and we reasoned that this might also result in increased bait avidity. In experiments to investigate alternative coupling methods, we observed that when we added a Cys residue to the C terminus of SC3, the purified SC3-Cys protein formed a dimer. We conjugated dimeric SC3 to beads using NHS chemistry, loaded these beads with the model bait mCherry-hFc-ST3 at varying concentrations, and then measured the amount of mCherry-hFc-ST3 loaded onto the beads by detection of mCherry fluorescence using flow cytometry or by measuring anti-FLAG signals on the FM3D. Using flow cytometry, we observed that signals with 10 nM mCherry on monomeric beads were near background levels, while 10 nM mCherry on dimeric beads produced a signal similar to that observed with 1000 nM mCherry on monomeric beads (Fig. 5A).

**Fig. 5.**
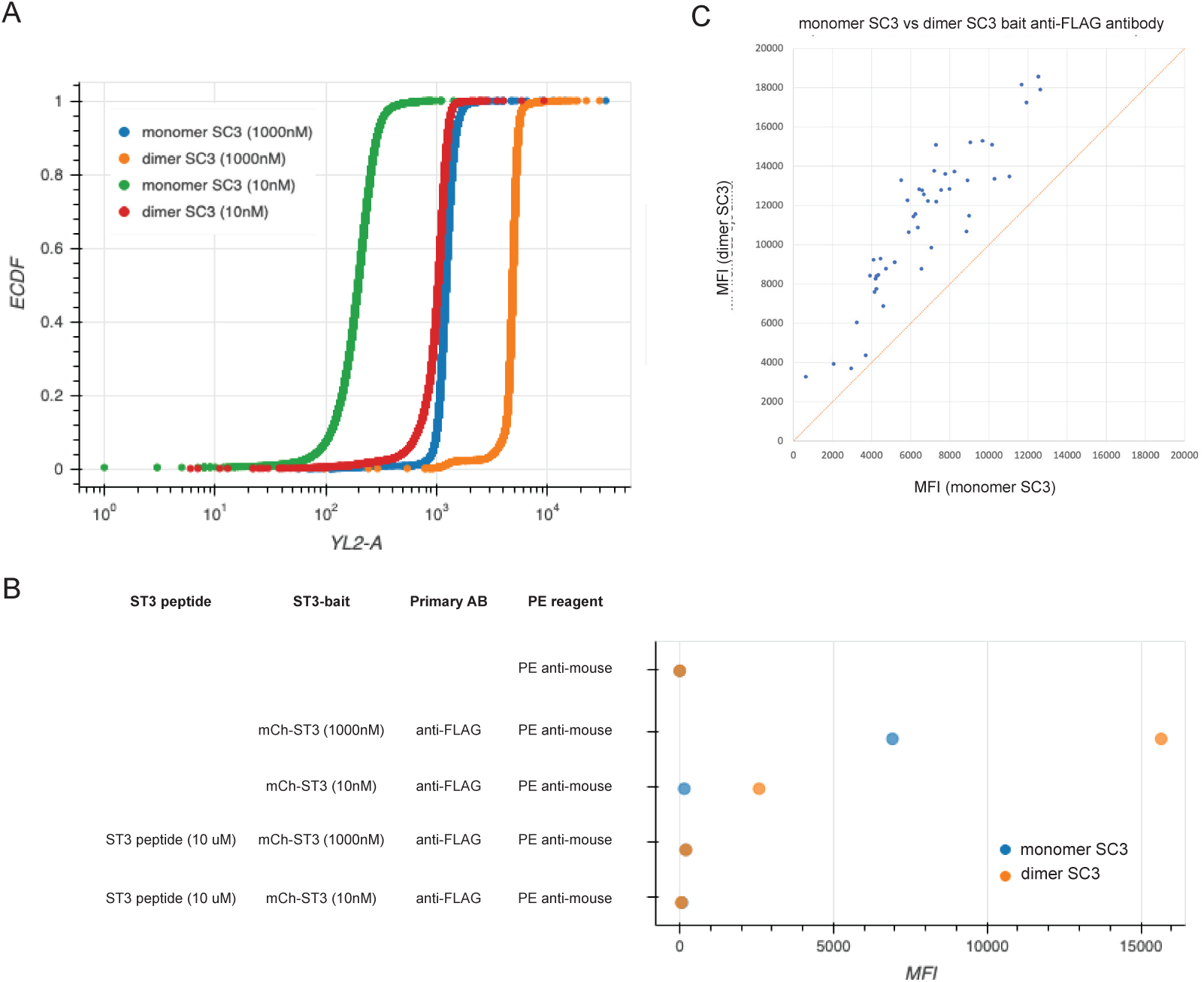
Using dimeric SC3 beads increases bait loading. A. Flow cytometry analysis of mCherry levels, using intrinsic mCherry fluorescence (YL2-A channel) on monomeric and dimeric SC3 beads loaded with mCherry-hFc-ST3 at 1000 nM or 10 nM concentrations. The signal for monomeric beads with 10 nM mCherry is at or near background levels. Note that the signal for dimeric beads with 10 nM mCherry is similar to that for monomeric beads with 1000 nM mCherry. The signal for dimeric beads with 1000 nM mCherry is ∼4-fold higher than with 10 nM mCherry. B. A graph of MFI values on the FM3D for mCherry-hFc-ST3 on monomeric and dimeric SC3 beads. Rows 1, 4, and 5 are negative controls. Row 2 (anti-FLAG, measures mCherry, on beads loaded with 1000 nM mCherry): MFIs for dimeric beads are ∼2.3-fold higher than for monomeric beads. Row 3 (anti-FLAG, on beads loaded with 10 nM mCherry): MFIs for dimeric beads are ∼7-fold lower than for dimeric beads loaded with 1000 nM mCherry. MFIs for monomeric beads are at background levels. Rows 4 and 5: blocking with ST3 peptide prior to adding mCherry-hFc-ST3 eliminates signals. C. Graph of anti-FLAG MFI values for all 41 PoC proteins loaded on monomeric (x-axis) and dimeric (y-axis) SC3 beads. Note that there is a consistent shift to higher MFI values with dimeric SC3 beads.

**Fig. 5, supplementary figure 1.**
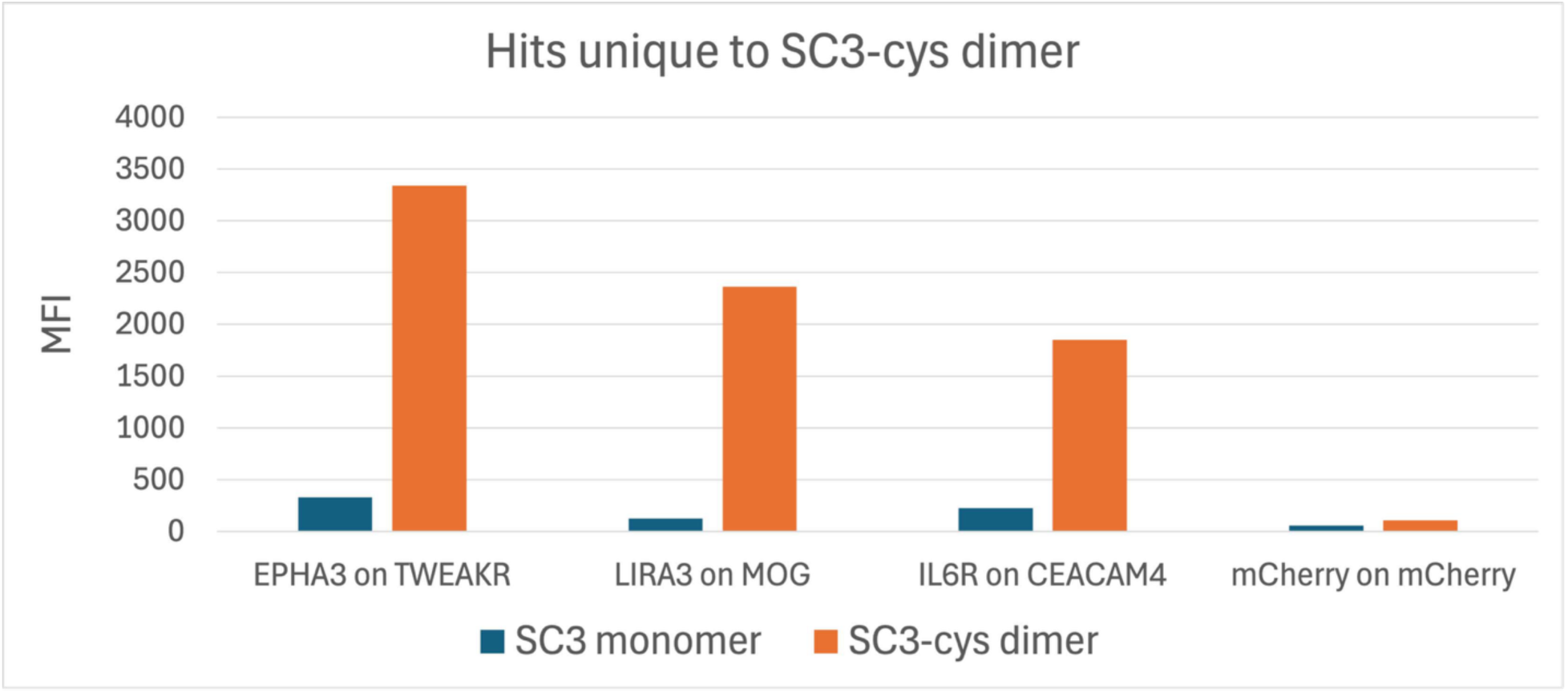
The bar graph shows MFIs for three hits in a run (M2/D1) in which preys were incubated with a bead region pool containing both monomeric SC3 and dimeric SC3 beads. We observed that these three interactions passed our statistical tests for dimeric SC3 beads, but had MFIs near background levels on monomeric SC3 beads. These are among the 15 new hits listed in Fig. 7, supplementary table 1. See Figs. 6 data supplement for raw data from these and other runs.

On the FM3D, MFI values for mCherry-hFc-ST3 loaded onto dimeric SC3 beads at 1000 nM were about 2.3-fold higher than for monomeric SC3 beads. At 10 nM mCherry, MFIs for monomeric beads loaded with mCherry at 10 nM were at background levels, similar to what we observed with flow cytometry. Dimeric beads loaded with 10 nM mCherry generated a strong signal (∼2500), although this was still ∼7-fold lower than the MFI signal observed with 1000 nM mCherry (Fig. 5B).

We loaded all 41 PoC set ECD-hFc-ST3 proteins onto dimeric and monomeric SC3 beads. We used different bead regions coupled to dimeric and monomeric SC3 and analyzed all of them together by anti-FLAG and anti-Fc staining (82 bead regions plus controls). Bait was coupled at the standard concentration (500 nM where possible). All MFI values were higher for dimeric SC3 beads (Fig. 5C).

**Figure 4, data supplement (a separate file)**

We then conducted screens with the 41 PoC preys against pools of the 41 PoC baits on dimeric and monomeric SC3 beads. All PoC set PPIs observed with the monomeric SC3 beads were also seen with the dimeric SC3 beads, and most PPIs had increased MFIs on the dimeric SC3 beads. However, PoC network PPIs (those found in either HIP1 or GNE) residing to the left of the knee on monomeric SC3 beads did not shift to the right side of the knee. Nevertheless, the dimeric SC3 provided a significant improvement to the sensitivity of the assay, which is important for ECD-hFc-ST3s expressed at low levels and for detection of PPIs with low signal strength. In fact, three candidate new PPIs were only seen when dimeric SC3 beads were used (Fig. 5, supplementary Fig. 1). The increase in signal strength on the dimeric SC3 beads might arise solely from the higher amounts of bait protein on the beads, or via a combination of the higher amounts of bait and increased bait avidity. A dimeric SC3 might couple to ST3s on two different ECD-hFc-ST3 dimers, such that one SC3 dimer is associated with four ECDs (Fig. 1B).

### Statistical analysis of binding interactions

We repeated the screen protocol described above with dimeric SC3 beads. In this paper, we primarily consider four runs, designated as D1-D4. D1 and D2 are technical replicates, run on the same day, while D3 and D4 were run on separate days. In addition, because there were a few expected hits not observed in one or more of these runs due to low protein concentration or low bead count, we repeated the screen with new preys and baits in another pair of runs, D5 and D6, and recovered these hits, as described below.

Fig. 6A is a graph of MFIs for D1 vs. D2, which shows that the observed values are highly reproducible between technical replicates. The data for these two runs are displayed in a histogram of binned MFI data in Fig. 6B. This is plotted on a log scale since the vast majority of data points have MFIs of 100 or less. The data are spread over a wide range of MFI values, with a minimum around 800-1000.

**Fig. 6.**
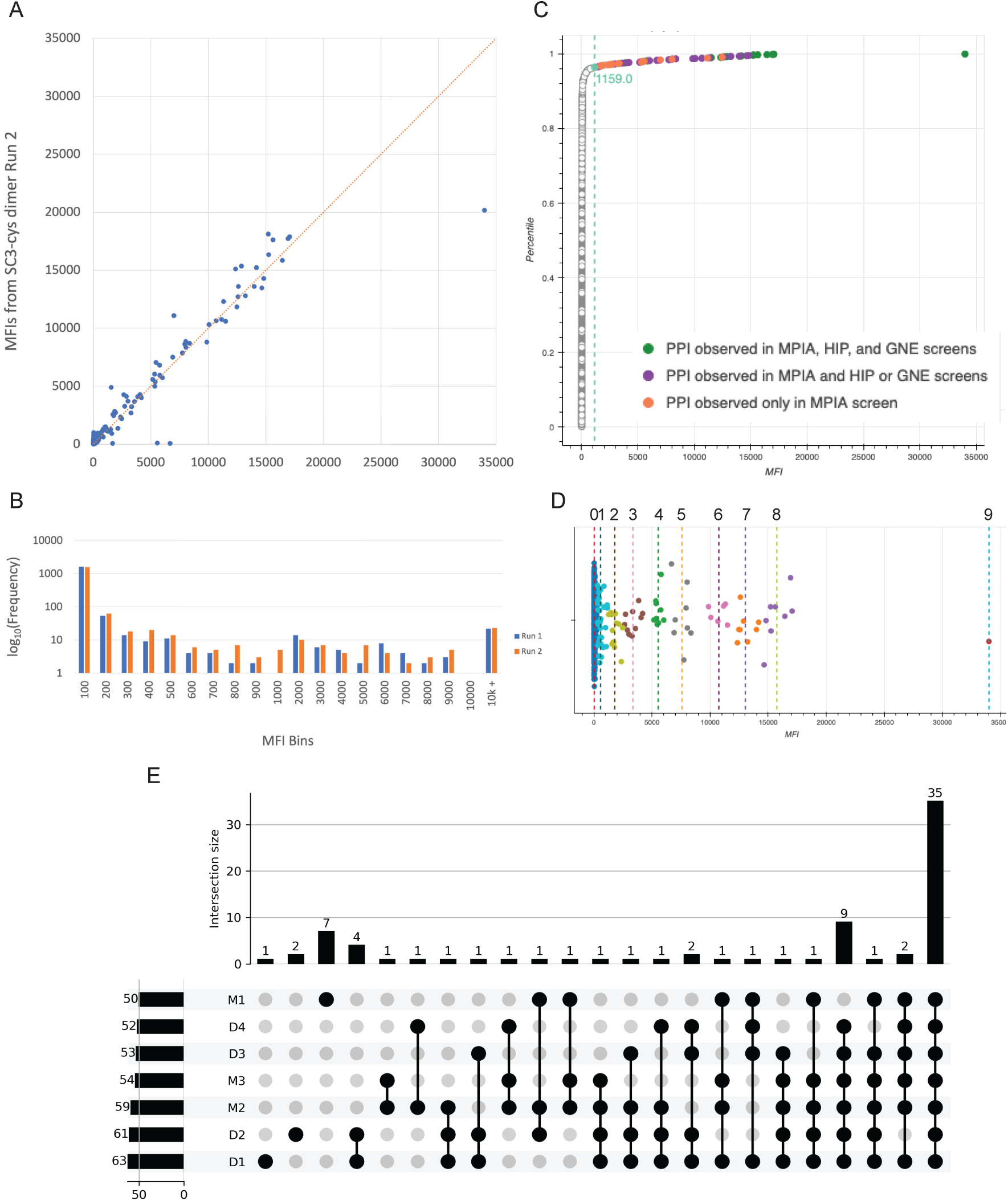
MPIA screen results and analysis of data. A. A graph of MFI values for all prey::bait pairs for run D1 (x-axis) vs. run D2 (y-axis). These are technical replicates, done on the same day with the same baits and preys. Note that the points are tightly clustered along the line. B. Histogram of binned MFI values for runs D1 (blue) and D2 (orange). The y-axis is plotted on a log scale, because the vast majority of data points have MFI<100. C. ECDF plot of run D1. The x-axis is MFI. y-axis is cumulative probability. See Fig. 4 legend for notes on ECDF plots. Green dots represent all hits found by MPIA, HIP1, and GNE, as in Fig. 4. Purple dots are hits observed by MPIA and by either HIP1 or GNE. Orange dots are new hits observed only by MPIA, which are listed in Fig. 7, table supplement 1. Note that all of these are to the right of the knee point, which is at MFI 1159 and is indicated by the teal dot and colored line. D. K-means clustering of data for run D1. The median MFIs for clusters 0-9 are indicated by dotted lines. Clusters 1-9 were considered to be possible hits and were analyzed using the outlier and ECDF filters to establish a hit list. E. Upset plot of overlap of hits selected by our statistical filters among dimer bead (D1-D4) and monomer bead (M1-M3) runs. Run M1, done earlier, used an entirely different set of proteins from the other runs. There is more divergence between run M1 and the remaining runs than between other groupings. The seven hits unique to M1 are probably false positives due to nonspecific ICAM5 prey binding to multiple bait beads. M1 ICAM5 prey only bound to a fraction of baits, so these hits were not eliminated by the outlier filter, but they are likely not real because one is for a negative control bead. The nine hits that were not found in M1 but were seen in all the other runs are probably explained by better expression of proteins used for the later runs. **Figure 6, data supplement (a separate file)**

**Figure 6, supplementary figure 1.**
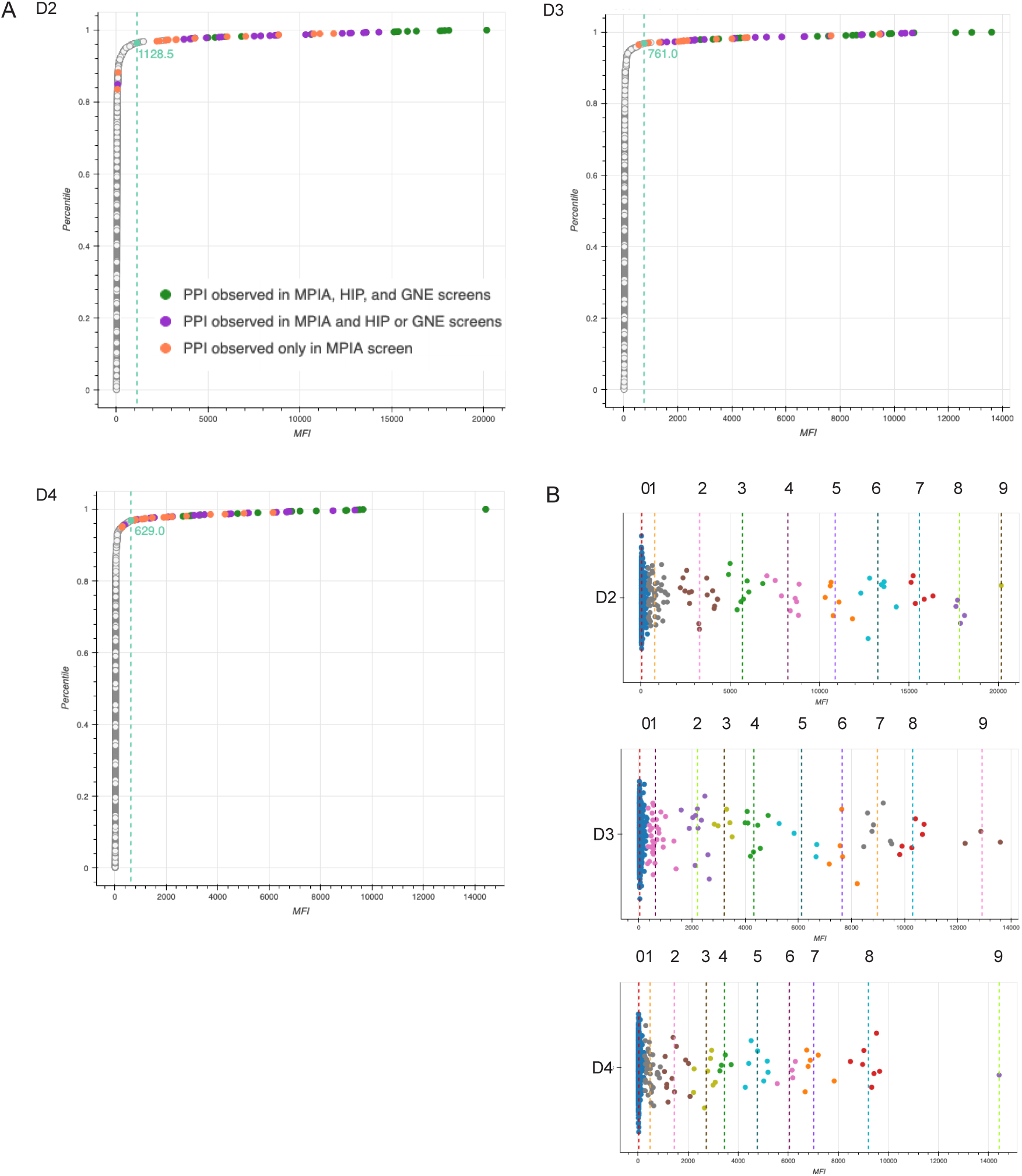
ECDFs and K-means plots for runs D2-D4. See Fig. 6 legend for details on these plots.

To assign hits from these runs, we devised a three-part data analysis framework. First, we employed K-means clustering, an unsupervised machine learning algorithm, to segment our data by MFI into distinct categorical groups that correspond to different levels of confidence with respect to our hits. The algorithm iteratively assigns each data point to its respective cluster’s nearest mean, and means are recalculated until the distances between the means and their respective data points are minimized, such that the means become fixed. The choice of number of clusters is dependent on the structure of the dataset. Since the strong hits are obvious, the utility of the K-means analysis is in identifying weaker potential PPIs that have MFIs in the border region where hits and non-hits are mixed. Given this criterion, it was important to choose a cluster number that segmented the data at a level of granularity such that samples that fell within the baseline noise level would be included for further analysis. For this set of screens, we found that choosing 10 clusters was appropriate. We designated cluster 0 as non-hits and funneled clusters 1-9 through the rest of our analytical pipeline (Fig. 6D).

With K-means clustering, we draw a relatively lax threshold for hits. Given this, we expect that some of the points in the lowest clusters result from nonspecific binding interactions that generate MFIs within the same range as the weak hits. There is also inter-experimental noise variability, since in some runs we observed “sticky” baits and preys that exhibit more noise than is typical due to nonspecific binding to many other proteins. To account for these factors, we devised a group-wise statistical outlier detection filter. Data were grouped by prey and bait, and for each group, the quartiles and the interquartile range (IQR) were computed. The outlier threshold was set to the maximum data point that is within 1.5 × 𝐼𝑄𝑅 above the 75^th^ percentile of the group. Data points seen above this threshold were considered to be outliers. Only prey::bait pairs identified as outliers relative to both other preys and other baits (in separate prey- and bait- oriented analyses) were considered to be possible hits. For run D1, this method filtered out 19 potential hits.

Lastly, we analyzed the ECDF of MFI values for each dataset. The ECDF was computed and sorted by MFI, and the knee points identified using the kneedle algorithm. This approach identifies the point of maximum curvature in the distribution, providing an orthogonal threshold (∼20-fold over median background) for hit identification. We then used this threshold as an additional filter to classify hits as those with MFIs above the knee. The knee is at an MFI of 1159 for D1 (Fig. 6C), and 1128 for D2. ECDFs and K-means plots for D2-D4 are in Fig. 6, supplementary figure 1. Raw data from D1-D4, as well as from three monomeric SC3 bead runs (M1-M3) are in Fig. 6 data supplement.

Using these algorithms, we identified a total of 54 hits that were seen in three of the four runs. Only 48 were seen in all four runs, but four of the six missing hits are accounted for by the fact that ICAM5 prey was not added in run D2 and bead count was below threshold for WFIKKN2 bait in run D4. We generated a plot of overlap (an upset plot) among the four runs, as well as three runs with monomeric SC3 beads (M1-M3), and this is shown in Fig. 6E.

### Benchmarking MPIA data against the HIP1 and GNE screens

Between them, the HIP1 and GNE screens identified 39 binding pairs in the PoC network. Nine were seen in both HIP1 and GNE, and the other 30 only in one of the screens. As in the monomeric SC3 bead screens described in Fig. 4, we identified all nine binding pairs found in both HIP1 and GNE in every run with dimeric SC3 beads (Fig. 7C). With the exception of BTNL9::IL6R (convention is prey::bait), these were strong hits in both prey::bait and bait::prey orientations, with MFIs ranging from 2117 to 33997. The BTNL9-IL6R pair was only detected when IL6R was the prey. Interestingly, this was also the case in the HIP1 and GNE screens.

**Fig. 7.**
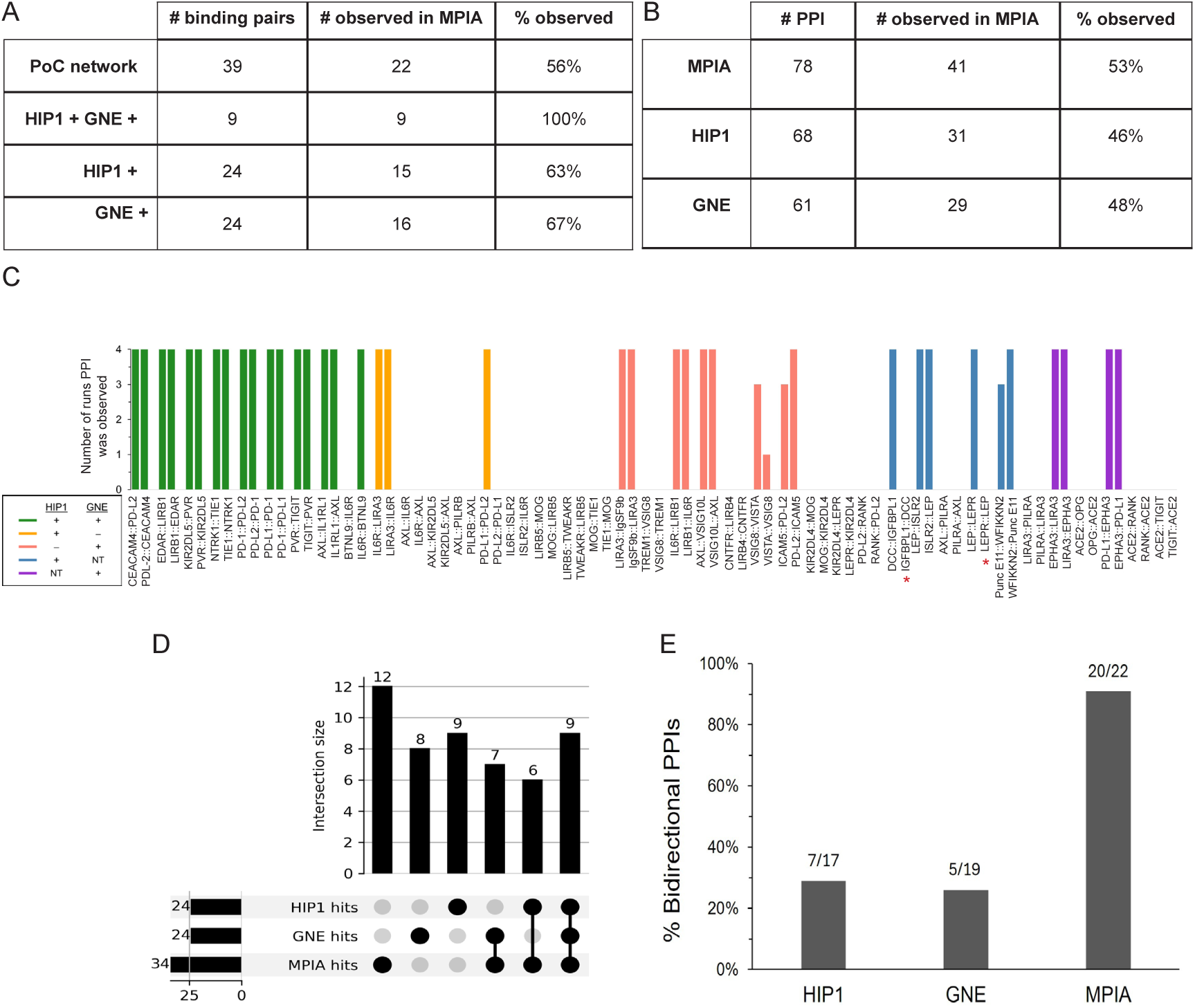
Comparison of MPIA data with HIP1 and GNE results. A. Table comparing recovery of PoC network binding pairs by the three screens. For the MPIA, we are defining hits as those observed in three of the four D1-D4 runs. B. Table comparing recovery of PoC network PPIs (two per binding pair) by the three screens. C. Bar graph showing PoC network PPIs observed as hits in MPIA runs D1-D4. The color indicates whether the PPI was found in HIP1, GNE, or both (see key). The x-axis shows the PPIs grouped by binding pairs. The y-axis shows the number of runs in which a PPI was observed. Asterisks denote two PPIs not observed in D1-D4 but found in other runs (see Results text). D. Upset plot and bar graph of overlap of binding pairs within the PoC protein set among MPIA, HIP1, and GNE screens. Seventeen binding pairs found in HIP1 or GNE were not seen in the MPIA, and 12 new binding pairs were observed in the MPIA that were not seen in either HIP1 or GNE, and therefore are not in the PoC network. The horizontal bars indicate the total number of binding pairs defined within the PoC protein set by each method. E. Graph of bidirectionality. The MPIA detects a much higher percentage of interactions bidirectionally than HIP1 or GNE. **Figure 7, data supplement (a separate file)**

**Fig. 7, supplementary Table 1.** New hits identified in MPIA screens with dimeric SC3 bait beads. MFIs are listed for each run in the last four columns. *Top*. PPIs for binding pairs that were not observed in HIP1 or GNE screens. PPIs with ICAM5 prey were not seen in run D2 because that prey was not added to the well as the result of an experimental error. *Bottom*. PPIs in a prey::bait orientation that was not observed in HIP1 or GNE screens. In the HIP1 or GNE screens, these PPIs were observed in the opposite prey::bait orientation. We also detected EPHA3::LIRA3 and EPHA3::PD-L1 as hits. In these cases, this orientation was not tested in either screen, but the reverse orientation was observed as a hit in GNE.

| Prey::Bait | # Runs Observed | Bidirectional | Run D1 | Run D2 | Run D3 | Run D4 |
| --- | --- | --- | --- | --- | --- | --- |
| EDAR::ICAM5 | 4 | yes | 2726 | 3277 | 2032 | 2016 |
| ICAM5::EDAR | 3 | yes | 1671 | 67 | 2230 | 2787 |
| EPHA3::LEP | 4 | yes | 11163 | 10762 | 7666 | 5024 |
| LEP::EPHA3 | 4 | yes | 12477 | 11838 | 9462 | 6155 |
| MOG::ICAM5 | 4 | yes | 5329 | 6041 | 4571 | 1459 |
| ICAM5::MOG | 3 | yes | 5563 | 100 | 4000 | 3722 |
| IL6R::TIGIT | 4 | no | 5449 | 7052 | 3429 | 2218 |
| EPHA3::TWEAKR | 4 | no | 3336 | 3235 | 2140 | 805 |
| IL6R::CEACAM4 | 4 | no | 1850 | 2830 | 1322 | 1059 |
| IL6R::IgSF9b | 4 | no | 8025 | 8849 | 4474 | 4286 |
| LIRA3::MOG | 3 | no | 2363 | 2354 | 920 | 300 |
| IL6R::IL6R | 4 | homophilic | 2645 | 4279 | 2121 | 1200 |
| TIE1::TIE1 | 4 | homophilic | 3280 | 2695 | 2355 | 1906 |
| LIRB4::LIRB4 | 3 | homophilic | 2469 | 2208 | 564 | 765 |
| VSIG8::VSIG8 | 3 | homophilic | 7000 | 11087 | 918 | 273 |

PPIs in prey::bait orientation not observed in HIP1 or GNE screens
| Prey::Bait | # Runs Observed | Bidirectional | Run D1 | Run D2 | Run D3 | Run D4 |
| --- | --- | --- | --- | --- | --- | --- |
| LIRA3::IL6R | 4 | yes | 10068 | 10313 | 6693 | 5194 |
| PD-L2::PD-L1 | 4 | no | 4197 | 4013 | 2206 | 1087 |
| IgSF9b::LIRA3 | 4 | yes | 1560 | 4906 | 2603 | 1402 |
| IL6R::LIRB1 | 4 | yes | 2885 | 4126 | 1899 | 1536 |
| AXL::VSIG10L | 4 | yes | 6017 | 5741 | 4077 | 3086 |
| VSIG8::VISTA | 4 | yes | 6906 | 7514 | 1404 | 377 |
| PD-L2::ICAM5 | 4 | yes | 8046 | 8362 | 4866 | 3270 |
| ISLR2::LEP | 4 | yes | 13992 | 13608 | 8823 | 6677 |
| WFIKKN2::Punc E11 | 4 | yes | 14815 | 14294.5 | 10404 | 8484 |

For the 30 binding pairs that were only observed in either the HIP1 screen or the GNE screen, but not both, we observed 13/30 (43%). Thus, in total, the MPIA identified 22 of the 39 previously reported binding pairs in the PoC network (56%)(Figs. 7A, D). For most of the 17 binding pairs seen in HIP1 or GNE that we did not identify as hits in the MPIA screen, MFI signals were at background levels. The absence of binding for some of these pairs is curious, because those same proteins were able to participate in other strong interactions that we did detect (see below for further discussion).

There are 78 potential PPIs in the PoC network (two per binding pair). 68 were tested in HIP1 and 31 observed (46%), while in the GNE screen 61 were tested and 29 observed (48%). Of the 54 hits seen in three of the four MPIA runs with dimeric SC3 beads, 39 are PoC network hits; that is, they had been found in either HIP1 or GNE (Fig. 7B). The remaining 15 are new hits involving interactions among PoC set proteins that were not seen in either HIP or GNE. In addition to the 39 PoC network hits seen in D1-D4, IGFBPL1::DCC was always identified as a strong hit in screens done with other bait bead sets, but we did not observe it in D1-D4 because the DCC bait bead count was below threshold. In two subsequent screens, D5 and D6, IGFBPL1::DCC generated MFI signals of 9737 and 5942. DCC::IGFBPL1 was a strong hit in all six screens. LEPR::LEP was not detected in D1-D4 (MFIs of 398-640) because LEPR was expressed at low levels (160 nM). We re-expressed LEPR, obtained a concentration of 790 nM, and used this LEPR prey in D5 and D6, in which it produced MFI signals of 19152 and 7930. Interestingly, there was less of an effect of LEPR concentration on the bead, because LEP::LEPR had MFI signals between 1361 and 3682 in D1-D4, and this increased to only 4567-5927 in D5 and D6. This is consistent with the observation that bait loading is less sensitive to protein concentration. In summary, then, the MPIA found 56 total hits, and 41 of the 78 PoC network PPIs (53%) that had been seen in either the HIP1 or GNE screens (Fig. 7B). All of these data are compiled in the Fig. 7 data supplement.

Detecting an interaction in both directions increases confidence that one has identified a bona fide binding pair. One important difference between the MPIA data and the HIP1 and GNE results is the number of binding pairs with PPIs observed bidirectionality. Of the 22 binding pairs detected in the MPIA, 20 were observed bidirectionally (91%). By comparison, 7/17 (29%) of tested binding pairs were observed bidirectionally in HIP1, and 5/19 in GNE (26%)(Fig. 7E).

There are 26 hits for PoC set proteins that were only seen in the MPIA. Eleven were the reverse orientation of PoC network PPIs that had been detected in HIP1 or GNE, and nine of these had been tested in either HIP1 or GNE and recorded as non-hits. An additional 15 hits that passed our statistical tests defined 12 new binding pairs that have not been previously reported, to our knowledge. Thus, the MPIA found 34 binding pairs for PoC set proteins, while HIP1 and GNE each found 24 (Fig. 7D). Three of the new pairs were detected bidirectionally, and four are homophilic (Fig. 7, supplementary table 1). Interestingly, for two of the pairs seen only unidirectionally (EPHA3::TWEAKR and IL6R::TIGIT), we also detected the reverse orientation as a hit in one or two of the four runs. Finally, three of the new hits were only observed with dimeric SC3 beads (Fig. 5, supplementary Fig. 1).

In summary, the fact that we found 56 hits in our screens of PoC set proteins, while HIP1 found 31 and GNE found 29, is consistent with the idea that the MPIA is more sensitive due to its greater dynamic range and to the higher avidity of the SC3-mi3 prey particles relative to the pentamers used in standard ELISA-based methods. Dimeric SC3 on the beads may also result in higher bait avidity.

## Discussion

### The MPIA method

The generation of a human cell-surface interactome map is a daunting task. Because cell-surface interactions cannot be accurately assessed using methods such as Y2H or AP-MS (Shilts and Wright 2024), methods specific to ECD interactions are required. We believe that the multiplex screening method described in this paper, the MPIA, together with automation advances we have made that will be described separately, is the first to have the capacity to allow a small group to execute a global in vitro interactome screen of human cell-surface and secreted proteins.

In the MPIA, we can use ECD-hFc-ST3 proteins as both bait and prey because they are covalently attached to both the bait bead and the 60-mer mi3 nanoparticle via SC3, and the prey nanoparticle contains a V5 epitope-tagged adapter for detection of prey binding. Approximately 20% of the SC3 sites on SC3-mi3 nanoparticles are filled with the adapter and the remaining sites with ECD-hFc-ST3 (Fig. 1). As baits, the same fusion proteins are coupled to SC3-conjugated Luminex MagPlex^TM^ fluorescent beads. There are 500 colors (regions) of these beads, and they can be distinguished using a laser on the FM3D instrument. This allows us to mix up to 500 baits in a single well, add one SC3-mi3-adapter-ECD prey, and then read out prey binding and bait identity on the FM3D. The FM3D can screen a 384-well plate of beads, representing 384 preys screened against 500 baits, in a few hours (i.e., 192,000 prey::bait pairs screened for interaction in a single plate).

Although the MPIA was developed to screen interactions between proteins located in the extracellular space, the method could also be used to screen interactions among intracellular proteins. Additionally, a library of CSP and secreted bait beads, such as the one we are making that will comprise at least 2200 proteins, can be used to rapidly screen antibodies for off-target binding.

### The cell-surface interactome problem

Methods in which ECDs are expressed as soluble preys and baits and tested for binding in vitro can be applied to most cell-surface and secreted proteins. We estimate that there are about 1600 human genes encoding STM and GPI-linked proteins that are entirely or partially localized to the cell surface and have ECDs that can be expressed as soluble fusion proteins. The extracellular regions of most multispan transmembrane (MTM) cell surface proteins are composed of discontinuous loops, but a few hundred MTMs have discrete ECD domains that could be expressed in a soluble form. There are about 1800 genes encoding secreted proteins, several hundred of which encode ligands that bind to signaling receptors. Of course, the actual numbers of CSPs and secreted proteins that exist in humans are severalfold larger than these numbers, because most genes encode multiple isoforms, and additional isoforms are generated by proteolytic cleavage or covalent modification. Since the number of interactions that must be screened expands as the square of the number of proteins, the problem becomes much larger if multiple isoforms were to be expressed and screened. We and others who have conducted ELISA-based screens have usually expressed only one isoform per gene. The high throughput nature of the MPIA, which allows multiplexing of 500 baits in a single well, will make it possible for investigators to include more than one isoform in interactome screens, at least for selected protein families.

What fraction of the human cell-surface interactome has been analyzed thus far? The three ELISA screens discussed here (Shilts et al. 2022; Verschueren et al. 2020; Wojtowicz et al. 2020) have cumulatively examined <500,000 pairwise interactions, which is <10% of the total number of interactions that could be assessed in vitro (Shilts and Wright 2024). The interactome of secreted protein binding to signaling receptors is more complete. Most of the genes for secreted ligands encode proteins with least one identified receptor. However, we think that at least 200 secreted ligands are likely to have additional receptors that have not yet been identified and are therefore important to include in a global interactome screen (Siepe et al. 2022). In our screen, we plan to express all of these, together with the ∼1600 STM ECDs and the MTM ECDs that can be expressed as fusion proteins, for a total of >2200 proteins.

How many interactions are likely to be identified in such a screen? In the HIP1 screen, 564 proteins were screened (318,096 interactions) and 426 binding pairs identified. Similar numbers were obtained in the GNE and Shilts et al. screens. However, there is only ∼30% overlap among the hits identified for proteins that were examined in any pair of screens. This low concordance could be explained by hypothesizing that most hits are false positives, but this is not likely to be the case, since for all three screens a selected set of interactions were tested using one or more orthogonal assays, such as SPR and cell binding, and ∼90% were successfully validated (Shilts et al. 2022; Verschueren et al. 2020; Wojtowicz et al. 2020). If each screen identified 30% of interactions and there are few false positives, the total number of actual interactions among the 564 HIP1 ECDs is likely to be >1200, and for 2200 proteins it may be >5000.

### Comparisons with ELISA-based screens

Since the HIP1 and GNE screens were performed using almost identical methods, with ECD-hFc dimers on ELISA plates as bait and pentameric ECD fusions formed using the COMP domain as prey (AP fusion in HIP1 and beta-lactamase fusion in GNE), if the screens have a low false positive rate it is surprising that they had only 30% overlap. We thought that the most likely explanation for the low concordance was that interactions were not detected for some proteins that were expressed poorly, and the poorly expressed protein subset might be different for the two groups. Unpurified supernatants were used in both screens (Wojtowicz et al. 2020; Verschueren et al. 2020). The MPIA screen method alleviated this problem to some extent, because we purified all ECD-hFc-ST3 proteins, which concentrated them by ∼6-fold relative to supernatant, and we conducted QC experiments on the FM3D to show that all preys and baits could generate signals.

In the MPIA screens described here, we examined a PoC set of 41 proteins (Fig. 2) that had also been screened in HIP1 and GNE. Together, those screens identified 39 protein binding pairs and 78 PPIs for these 41 proteins, and these 39 pairs define the PoC network against which we are benchmarking the MPIA screen. With the MPIA, we found all nine of the protein binding pairs found by both HIP1 and GNE, as well as 13 pairs that were seen in either HIP1 or GNE, but not both. ∼90% of binding pairs seen in the MPIA were detected bidirectionally, while pairs were detected bidirectionally in only ∼20% of cases in HIP1 and GNE (Fig. 7E). Detecting an interaction bidirectionally increases confidence that it is a true positive. Bidirectional detection is primarily responsible for the fact that we found 41 PoC network PPIs in the MPIA, vs. 31 and 29 in HIP1 and GNE (Fig. 7B).

Seventeen binding pairs in the PoC network that were only reported in one of the two previous screens were not detected by the MPIA (Figs. 6, 7). Seven involved proteins expressed at low levels, and these may have generated signals that were too low to pass our statistical tests. However, there are some interesting issues raised by examining the 10 interactions involving proteins expressed at moderate to high levels that we did not find with the MPIA. Several of these missed hits involved proteins that were capable of binding as preys and baits. For example, KIR2DL5 prey bound to PVR bait (MFI 5355) and ILRL1 prey bound to AXL bait (MFI 5785), but KIR2DL5 prey failed to bind to AXL bait (MFI 31), although KIR2DL5::AXL was observed as a hit in HIP1 (Fig. 7, data supplement). One possible explanation for this is that one of these proteins has a glycan at a site(s) that blocks its binding to one ligand but not to another, and that this was not present on the AP5 prey and/or Fc bait protein made in HEK293F cells in the HIP1 screen. It might have been present in the GNE protein(s), however, since they did not detect KIR2DL5 binding to AXL. To evaluate this hypothesis, we deglycosylated our ECD-hFc-ST3 proteins using N and O-glycosidase mixes (New England Biolabs) and ran a PoC set screen. However, we observed that this weakened most interactions and did not uncover KIR2DL5::AXL or other missing HIP1/GNE hits (data not shown). Perhaps carbohydrates that are missing from deglycosylated KIR2DL5 and/or AXL are required for their binding, so complete deglycosylation does not generate proteins that are capable of binding. Alternatively, the differences between the screens might be due to other factors.

In HIP1 we used phylogenetic homology analysis (PHA) to predict false negatives in the ELISA-based binding screen and tested them for binding by SPR. Our rationale for doing this was that phylogenetically related subfamily members often exhibit binding to one another. We reasoned that, in cases where we observed binding of one subfamily member to a particular partner (protein X) but not binding of other related subfamily members to protein X, these missing interactions may have been false negatives. Consistent with this reasoning, when we performed SPR experiments on several different subfamilies of non-binding phylogenetically related members, we found that 33/35 of the pairs that did not bind in the ELISA screen did bind on an SPR chip (“SPR-only” PPIs). This was curious because the ligand and analyte proteins were purified versions of the same ECD-hFc used in the ELISA-based screen. This suggested that the dimeric ECD-hFc fusion proteins might be able to bind to one another but not to the pentameric ECD-AP5 proteins.

One example of this is IGFBPL1, which bound to DCC in the HIP1 screen but not to other members of the DCC subfamily, which includes PuncE11. However, when the five DCC subfamily members were tested for binding to IGFBPL1 by SPR, all five were found to bind, including PuncE11. WFIKKN2, a protein from a different subfamily from IGFBPL1, bound to PuncE11 in the HIP1 screen but not to DCC. However, WFIKKN2 did bind to DCC when tested by SPR (Wojtowicz et al. 2020). We included all four of these proteins in our PoC set for the purpose of evaluating whether HIP1 SPR-only PPIs would bind in the MPIA. We predicted that they might because, in the MPIA, both prey and bait are dimeric hFc. In our MPIA screen, IGFBPL1 bound to DCC, and WFIKKN2 bound to Punc E11. These were strong interactions (MFI >7000) in both orientations (Fig. 7, data supplement). However, as in the HIP1 screen, we did not detect any binding of IGFBPL1 to PuncE11, or of WFIKKN2 to DCC, although all of these proteins were clearly capable of binding in our screen. This suggests that there are differences between the interactome and SPR assays that affect whether proteins can bind. One difference is in how the bait protein is affixed to the surface to which it is coupled. In ELISA-based assays and the MPIA, bait protein is coupled to plate and bead surfaces, respectively. By contrast, in SPR, bait protein (called ligand) is attached to the chip surface via a long flexible dextran linker. As such, it has freedom of movement that resembles a protein in solution. To try to approximate these conditions, we made beads in which ECD-hFc-ST3 baits were linked to the bead surface by a flexible 10 kD PEG linker. However, we did not find HIP1 SPR-only PPIs in experiments conducted with these beads (data not shown), suggesting there are unknown biophysical differences between the assays that we have not been able to address.

We identified 12 new binding pairs within the PoC set that were not published or seen in previous screens, three of which were detected bidirectionally (Fig. 7, supplementary table 1). Validation of these pairs using other methods such as SPR and cell binding will be the focus of subsequent studies. Assuming that we are able to validate these new interactions, the MPIA screen appears to be somewhat more sensitive than ELISA-based screens, since we found a total of 34 binding pairs in the PoC set, while HIP1 and GNE each identified 24 (Fig. 7). We plan to now screen the entire set of 564 HIP1 proteins using the MPIA, retest all significant candidate interactions, and then evaluate overlap among the screens with this larger set. We also plan to use PHA to select additional proteins from interacting subfamilies to test by medium-throughput assays such as cell binding and multiplex SPR. In this manner, we hope to define a network that more closely resembles the complete interaction map than that defined by any previous screen.

## Materials and Methods

### Molecular Cloning

The vector used for cloning PoC set ECDs for MPIA screening was derived from plasmids used in HIP1. It has a CMV promoter, HA SP, a Not I site, mCherry, an Asc I site, human Fc with the DAPA mutations, a FLAG tag, an AviTag, SpyTag003, and an 8X His tag (see Fig. 1). ECD sequences were transferred from the plasmid used in HIP1, in which the ECD region is also flanked by Not I and Asc I sites, into the MPIA vector, replacing mCherry. Sequences of the plasmid constructs (backbone and ECD) are in Fig. 2, data supplement, sheet 2.

The plasmid encoding N-terminally His-tagged SpyCatcher003 was obtained from Addgene (Cat# 133447). The plasmid encoding N-terminally His-tagged SpyCatcher with a C-terminal cysteine was developed in-house via ligation of a EcoRI- and MfeI-digested synthetic dsDNA fragment encoding the C-terminal cysteine (IDT gBlock) into the gel-extracted EcoRI- and MfeI-digested Addgene plasmid backbone. The plasmid encoding SpyCatcher003-mi3 with a C-terminal C-tag was obtained from Addgene (Cat# 159995).

### Preparation of transfection-quality DNA

We used the Macherey-Nagel NucleoBond Xtra Maxi Plus Maxiprep (#740416) Protocol:

Day 0

1. Prepare a 4mL preculture in LB with kanamycin

a. 250RPM/37C

Day 1

1. Inoculate 4mL preculture in 120mL Plasmid+ media with kanamycin

a. 250RPM/37C

Day 2

1. Harvest pellet at 6000g/15minutes/4C

a. Dry pellet as much as possible
2. Resuspend pellet in 24mL Resuspension buffer
3. Add 24mL Lysis buffer and mix gently

a. Let sit for 5 minutes
4. Add 24mL Neutralization buffer and mix until mixture turns white
5. Equilibrate NucleoBond Xtra column with 25mL Equilibration buffer
6. Pour lysate into NucleoBond column
7. After lysate goes through the column, wash with 15mL Equilibration buffer around the rim of the column
8. Remove and discard column filter
9. Add 25mL Wash buffer to column
10. Place a 50mL centrifuge tube under the column and add 15mL Elution buffer
11. Add 10.5mL isopropanol to the eluate and vortex
12. Place a finalizer on a syringe and push lysate through
13. Add 4mL 70% ethanol and push through
14. Push air through syringe 15-20 times until dry
15. Aspirate 1mL TRIS buffer and attach finalizer
16. Push 0.5mL through finalizer and let sit for 5 minutes

a. Push final 0.5mL through finalizer and push air through to flush out the entire 1mL of TRIS

### Transfecting Expi293 cells using the Expifectamine 293 Transfection Protocol

We used 30 ml cultures for our experiments. Protocol:

Day 0

1. Expi293 cells are maintained at a density of 0.3-6e6

Day 1

1. Take a cell count and dilute cells to a density of 3e6 with prewarmed Expi293 medium

a. CO2: 8%
b. Temperature: 37C
c. Shaker speed:

i. 50ml Centrifuge Tubes: 470RPM
ii. ii. Erlenmeyer Flasks: 130RPM
2. Prepare 2x Opti-mem tubes per transfection labelled DNA and EXPI

a. 50ul. Opti-mem per 1ml transfection
3. Add 1ug. /ml transfection of plasmid DNA to DNA Opti-mem tube and mix

a. Sterile filter plasmid DNA if necessary
4. Using a 3-minute timer, add 2.7ul. Expifectamine/ml transfection to EXPI Opti-mem tube and mix well
5. After 3-minutes are up, set a 5-minute timer and combine the DNA and EXPI Opti-mem tubes, mix well
6. After the 5-minutes are up, add the transfection mix to your 3e6 Expi293 cell culture (30 ml)

Day 2

1. Make enhancer mix in a tube (make a small overage, ∼20ul. extra):

a. Enhancer #1: Add 5ul. Per ml of transfection
b. Enhancer #2: Add 50ul. Per ml of transfection
2. Add enhancer to Expi293 cell cultures

Day 4-5

1. Harvest transfections @4000RPM/15min/4C

a. Harvest @6000RPM/15min/4C for 1-liter expressions
2. Filter supernatant (0.22um)

Materials

- Gibco Opti-MEM #31985062
- Expifectamine 293 Transfection Kit #A14525
- Expi293 Expression Medium #A1435101

### Purification of Proteins using the HisTrapFF Purification Protocol

We purified proteins from 30 ml of filtered supernatant to generate final purified preparations of 5 ml.

1. Hook up 1mL HisTrapFF column to the peristatic pump while running water at 1mL/minute.
2. Run 2 column volumes of water at 1mL/minute.
3. Equilibrate with 5 column volumes of wash buffer at 1mL/minute.
4. Load supernatant at 0.25-0.5mL/minute.
5. Wash 10 column volumes of wash buffer at 1mL/minute.
6. Elute 5 column volumes with elution buffer at 0.25-0.5mL/minute.
7. Measure protein concentration and analyze using SDS-PAGE, running 1 µl-15 µl of protein per lane, depending on expression level.

Buffers:

- Wash buffer: 1XPBS, 20mM imidazole, pH 7.5
- Elution buffer: 1XPBS, 500mM imidazole, pH 7.5

### Bead coupling to SpyCatcher003 and control NBs and antibodies

Approximately 500,000 Luminex MagPlex microspheres (aka “beads) for each bead region were coupled to SpyCatcher003 or SpyCatcher003-cys using NHS chemistry in 1.5 mL tubes. All aspiration steps were carried out in an Invitrogen DynaMag-2 magnetic rack (Cat# 12321D).

Briefly, beads were washed with activation buffer (0.1M NaH_2_PO_4_, pH 6.2) twice and resuspended to 80uL activation buffer. 10 µL each of sulfo-NHS (Thermo Cat# 24510) and EDC (Thermo Cat# 22980) at 50mg/mL in activation buffer were added to the beads and the reaction was incubated for 20 min at room temperature. Beads were then washed twice with 1x PBS and purified SpyCatcher003 or SpyCatcher003-cys was added to a final concentration of 25 ng/uL in 1x PBS to a final volume of 500 µL. The coupling reaction was allowed to incubate at room temperature for 2 hr with occasional mixing. Following coupling, the beads were washed twice with 1x PBS and then resuspended into 500 µL of 1% casein in 1x PBS (Thermo Cat# 37528) for blocking and storage.

An anti-GFP nanobody (His-14-AviTag-SUMOStar-anti-GFP nanobody 3K1K) (gift from Tino Pleiner) and a polyclonal goat anti-human Fc gamma antibody (Jackson Cat# 109-005- 098) were coupled to beads as above, to different bead regions.

### ECD-hFc-ST3 bait immobilization onto SC3-coupled magnetic beads

Purified ECD-hFc-ST3 bait proteins were normalized to 500 nM and 500 µl of protein was added to casein-blocked, SpyCatcher003-coupled beads. In cases where the purified protein concentration was less than 500 nM, the protein was used at its maximum available concentration. Each bait protein was added to a unique bead region in order to assign one region per bait protein. The beads were incubated with baits for 2 hr at room temperature with occasional mixing. Following the bait immobilization, beads were washed twice with 500 µl of 1% casein in 1x PBS. The beads were then resuspended into 100 µl of 1% casein in 1x PBS and then pooled to create the multiplexed bait bead set. The beads were then immobilized on a magnet and the supernatant was removed. The beads were then resuspended in 200 µl of 1% casein in 1x PBS containing SpyTag003 peptide (Bio-Rad Cat# BLP086) at 10 nM. Following SpyTag003 peptide blocking, the beads were washed twice with 500 µl of 1% casein in 1x PBS and stored at 4°C for later use.

### Making 60-mer SC3-mi3 prey

To create the adapter-coupled nanoparticle, GFP-2xV5-HA-ST3-8xHis adapter protein was mixed with purified SC3-mi3 at a ratio of 0.2 mol of GFP-2xV5-HA-ST3-8xHis adapter to 1.0 mol of SC3-mi3. Following overnight incubation at room temperature, a sample of the adapter- coupled SC3-mi3 was analyzed by SDS-PAGE to confirm the extent of coupling and to evaluate the concentration of the adapter-coupled SC3-mi3 and the remaining uncoupled SC3-mi3. ECD- ST3 was then added to the nanoparticle preparation at a ratio of 1.5 (ECD-hFc-ST3 monomer) to 1 (SC3-mi3-adapter) over the remaining vacant SC3 sites, to a final concentration of 1 µM of ECD-ST3 on nanoparticle. For preys with concentrations below 1uM, adapter-coupled SC3-mi3 was added to maintain a 1.5-2 molar ratio of ECD-hFc-ST3 monomer to SC3-mi3-adapter. For preys with concentrations below the detection limit of either Bradford assay or densitometry, adapter-coupled SC3-mi3 was added for an amount corresponding to ∼0.1uM of ECD-hFc-ST3. After overnight incubation at room temperature, selected samples were analyzed by SDS-PAGE to confirm ECD-hFc-ST3 coupling onto the SC3-mi3 nanoparticle.

### Multiplexed Protein Interaction Assay (MPIA)

The multiplexed, bait-loaded bead set was aliquoted into 96-well hardshell PCR plates (Bio-Rad) at a density of 500-1000 beads per region per well. All plate aspiration steps were performed by placing the 96-well plates onto a 96-well ring magnet (Alpaqua Cat# A001322) and aspirated using a Tecan Evo2 liquid handler. 20 µl of 60-mer preys, diluted to 333 nM ECD using 1% casein in 1x PBS, were added to the wells and the beads were gently resuspended 2- 3 times. After 2 hr incubation at room temperature, the plate was aspirated and then 50 µl of BioLegend mouse anti-V5-tag (Cat# 680602) at 0.5 ug/mL in 1% casein in 1x PBS was added to each well and resuspended. After 1 hr incubation at room temperature, the wells were aspirated under magnet and then 50 µl of BioLegend PE goat anti-mouse IgG (Cat# 405307) at 0.5 ug/mL was added to each well and resuspended. After 1hr incubation at room temperature, the plate was aspirated under magnet and then 100 µl of 1x PBS was added to each well. The beads were resuspended and then transferred to wells of a 384-deep well plate pre-filled with 100 µl of 1x PBS in each well. The 384-well plate was then loaded onto a Luminex FlexMAP 3D running in normal PMT mode and set to count 50 beads of each region defined in the assay for each well.

## Acknowledgments

This work was funded by an NIH TRO1 grant, GM150125, to K.Z., K. Christopher Garcia (Stanford), and Matthew Thomson (Caltech). Maxine Wang was supported by a fellowship from the H.S. Chau Foundation. Work at the Caltech Protein Expression Center is also supported by the Beckman Institute at Caltech.

Conceptualization, K.Z., M.A.A., W.M.W., J.V., M.L.W.; Investigation, M.A.A., M.L.W., E.G., A.W.L., P.V.; Data Curation and Visualization: W.M.W., M.A.A., M.L.W., K.Z.; Writing and Editing, K.Z., W.M.W., M.A.A., M.L.W.; Supervision, K.Z., W.M.W., J.V.

We thank Matt Thomson, Chris Garcia, Pamela Bjorkman, Barbara Wold, An Zhang, Verona Yue, Dirk Siepe, and Mitchell Guttman for helpful discussions, and Tino Pleiner for the anti-GFP NB construct.

## Notes

### Competing Interest Statement

The authors have declared no competing interest.

